# A chromosome-scale assembly for tetraploid sour cherry (*Prunus cerasus* L.) ‘Montmorency’ identifies three distinct ancestral *Prunus* genomes

**DOI:** 10.1101/2023.01.10.523438

**Authors:** Charity Z. Goeckeritz, Kathleen E. Rhoades, Kevin L. Childs, Amy F. Iezzoni, Robert VanBuren, Courtney A. Hollender

## Abstract

**Background:** Sour cherry (*Prunus cerasus* L.) is a valuable fruit crop in the Rosaceae family and a hybrid between progenitors most closely related to extant *P. fruticosa* (ground cherry) and *P. avium* (sweet cherry). Sour cherry is an allotetraploid with few genomic resources, so a genome sequence would greatly facilitate the improvement of this crop. In *Prunus*, two known classes of genes are of particular importance to breeding strategies: the self-incompatibility loci (*S*-alleles), which determine compatible crosses and are critically important for successful fertilization and fruit set, and the Dormancy Associated MADS-box genes (DAMs), which strongly affect dormancy transitions and flowering time.

**Results:** Here we report a chromosome-scale genome assembly for sour cherry cultivar ‘Montmorency’, the predominant sour cherry cultivar grown in the U.S. We also generated a draft assembly of *P. fruticosa* to use alongside a published *P. avium* sequence for syntelog-based subgenome assignments for ‘Montmorency’. Using hierarchal k-mer clustering and phylogenomics, we provide compelling evidence this allotetraploid is trigenomic, containing two distinct subgenomes inherited from a *P. fruticosa-like* ancestor (A and A’) and two copies of the same subgenome inherited from a *P. avium-like* ancestor (BB). We therefore assigned the genome composition of ‘Montmorency’ to be AA’BB and show little to no recombination has occurred between progenitor subgenomes (A/A’ and B). The *S*-alleles and DAMs in ‘Montmorency’ and *P. fruticosa* were manually annotated and demonstrated to support the three subgenome assignments. Lastly, the hybridization event that ‘Montmorency’ is descended from was estimated to have occurred less than 1.61 million years ago, making sour cherry a relatively recent allotetraploid.

**Conclusions:** The genome of sour cherry cultivar Montmorency highlights the evolutionary complexity of the genus *Prunus*. These genomic resources will inform future breeding strategies for sour cherry, comparative genomics in the Rosaceae, and questions regarding neopolyploidy.

## Background

Sour cherry (*Prunus cerasus* L.) is an important temperate tree crop whose fruit is valued for its uniquely sweet and acidic flavor and superior processing characteristics for products such as jam, juice, compote, and pie. Sour cherry is a member of the economically important Rosaceae family, which includes other cultivated *Prunus* species such as peach, sweet cherry, apricot, almond, and plum, as well as apples, pears, roses, strawberries, and various cane fruits (1,2). The evolutionary history of *Prunus* has been historically difficult to resolve as hybridization, polyploidy, and incomplete lineage sorting is rampant throughout the genus (1,3). Sour cherry is an allotetraploid (2n=4x=32) and shown to be a hybrid between a tetraploid resembling *P. fruticosa* Pall. (ground cherry) and a diploid resembling *P. avium* L. (sweet cherry) (4–8). However, cytological and genetic data suggest that sour cherry could be considered a segmental allotetraploid (5,9,10). Trivalents and quadrivalents are common at meiosis, and although disomic inheritance is more common, tetrasomic inheritance has also been observed (5,9–11). The native distributions for both *P. fruticosa* and *P. avium* overlap in central and eastern Europe and sour cherry exhibits intermediate phenotypes between these progenitor species (4,7,11,12). This hybridization event likely happened multiple times as *P. fruticosa, P. avium*, and *P. cerasus* all hybridize in the wild, and sour cherries with substantially different phenotypes occupy dissimilar hardiness zones (7,13). The *P. fruticosa-like* progenitor has been shown to be the more common maternal parent of sour cherry; however, some accessions resulted from the *P. avium-like* progenitor as the maternal parent (6,7,14). To add to the polyploid complexity, it is unclear if *P. fruticosa* is an allo- or autotetraploid as to our knowledge, no rigorous study has been reported (13,15–17).

Early spring freezes are major contributors to crop loss in the temperate fruit tree industry. For example, in 2012, an unseasonably warm March followed by an April freeze decimated tree fruit production throughout the midwestern United States (18). Michigan, the number one producer of sour cherry in the U.S., lost more than 90 percent of its crop (18). Depending on the region, climate change is increasing the frequency of these events worldwide (18–20). A better understanding of the genetic control of bloom time in fruit trees and breeding cultivars with later bloom times would reduce floral death and subsequent crop loss; as such, this has been a major goal for the sour cherry breeding program at Michigan State University (MSU). Sour cherries exhibit bloom times that span those of its two progenitor species, and it is hypothesized the alleles conferring later bloom time are derived from the *P. fruticosa-like* progenitor since *P. fruticosa* inhabits more northern latitudes compared to *P. avium* (21). Development of a sour cherry genome resource would support breeding efforts by enabling gene discovery and an understanding of the genetic basis of agronomic traits in this complex tetraploid. Until now, genetic studies have depended on traditional methods involving linkage maps and common markers and synteny between *Prunus* species (9,11,21), since no public sour cherry genome sequence is available.

In this work, we constructed and annotated the first *P. cerasus* reference genome for the cultivar ‘Montmorency,’ a ~400-year-old French amarelle sour cherry selection of unknown origin but the most widely-grown cultivar in the United States. We also sequenced, assembled, and annotated sequences from a *P. fruticosa* accession present in the MSU germplasm collection, which was used, along with a published *P. avium* genome (22), to assign subgenomes to the ‘Montmorency’ superscaffolds. To demonstrate the utility of the genome, we identified, manually annotated, and assigned progenitor subgenomes to two sets of genes present in the *Prunus* lineage. The first set includes the Dormancy-Associated MADS box genes (DAMs): highly conserved genes initially discovered in peach (*P. persica*) with major effects on dormancy transitions and flowering time in *Prunus* species (23–30). The second set includes the self-incompatibility *S*-allele genes that make up the *S*-haplotype, *S*-ribonuclease (*S*-RNase) and *S*-locus F-box (SFB) (31–41). The four *S*-haplotypes in ‘Montmorency’ have been previously characterized (38), but no *S*-haplotypes in *P. fruticosa* have been thoroughly described. Our findings for these two sets of genes are discussed in the context of their putative subgenome origin and possible influence on flowering time and floral self-compatibility.

## Results

### Assembly of the sour cherry ‘Montmorency’ genome reveals 3 distinct Prunus subgenomes

We generated a chromosome-scale reference genome for sour cherry ‘Montmorency’ using a combination of PacBio long-read and Illumina short-read sequencing for genome assembly and chromosome conformation capture (Hi-C) for scaffolding (**Supplementary Figure 1**). PacBio reads were assembled using Canu and polished with Pilon. The polished ‘Montmorency’ contigs have a total assembly size of 1066 Mb, or 172% of the estimated genome size of 621 Mb according to a k-mer analysis (k=25). The size of the assembly in conjunction with the abundant estimated heterozygosity (4.90%) suggests multiple haplotypes were assembled. The Merqury plot (42) (**Supplementary Figure 2**) indicates high genome completeness as most k-mers in the Illumina dataset are found in the assembly. Additionally, k-mers from the Illumina reads are found in the assembly at the expected relative frequencies and there are 4 distinct peaks, indicating haplotypes are well-phased. A BUSCO assessment (43) demonstrated the assembly contains a suitable representation of the gene space (>98% complete BUSCOs) and most of them are duplicated (>93%), as expected of a polyploid.

As sour cherry is an allotetraploid derived from two progenitor species, we expected to assemble two full haplotypes for ‘Montmorency’ (*Prunus* subgenomes; n=2x=16) with additional alleles being unanchored. To our surprise, initial scaffolding resulted in 24 linkage groups (chromosomes), 8 of which experienced sudden drops in signal along the Hi-C diagonal and were made up of significantly smaller contigs than the other 16. Preliminary phylogenomic analyses (see Methods) showed genes on these 8 poorly-scaffolded linkage groups were most likely derived from the *P. avium*-like ancestor. Therefore, we posited there was a disproportionate collapse of *P. avium-like* haplotypes compared to the others, with the poorer scaffolding being a result of collapsing and haplotype switching. From here, efforts were made to reassemble the genome with the goal of either 1) better phasing the *P. avium-like* sequences or 2) forcing them to collapse into a monoploid representation. Unfortunately, due to the heterogeneous nature of all four haplotypes’ sequences, the first scenario resulted in poor sequence correction and assembly inflation while the second negatively affected the assembly of the haplotypes that had previously been intact. Instead, Purge Haplotigs (44) was used to set aside one of the two possible alleles of the *P. avium-like* sequences. Since we knew these sequences consisted of smaller contigs, those greater than 400 kb that had been removed by Purge Haplotigs were added back into the assembly prior to scaffolding. **Supplementary Figure 3** suggested promising results: k-mers from the Illumina dataset were found only once in the purged (removed) assembly portion. In other words, few to no sequences common to multiple alleles (2x, 3x, 4x) were in this portion of the assembly.

The scaffolding pipeline was subsequently rerun, and 771.8 Mb (124% of the estimated haploid genome size) was scaffolded into 24 linkage groups (**Supplementary Figure 4**). These chromosomes were numbered according to their synteny with *Prunus persica* chromosomes 1 – 8 (**Figure 1a**) (45,46). These results and the Hi-C matrix suggested three homoeologs, instead of the predicted two, were assembled for each *Prunus* ancestral linkage group. We subsequently named and clustered these 24 chromosomes into three groups of 8 (A, A’, and B) based on their 25-mer signatures and progenitor assignments (details to follow). A comparison between the 24 linkage groups and the recently-published *P. avium* ‘Tieton’ genome (22) indicated substantial synteny amongst the ‘Montmorency’ chromosomes themselves as well as *P. avium* (**Figure 1b**), in concordance with numerous reports that *Prunus* genomes are highly syntenic (47–49). When the scaffolded linkage groups were compared with a sour cherry genetic map of 545 unique markers (11), 78% mapped exactly once to each homoeolog. Furthermore, all markers showed nearly perfect linearity with the assembly (**Figure 2, Supplementary Figure 5**).

**Figure 1:**
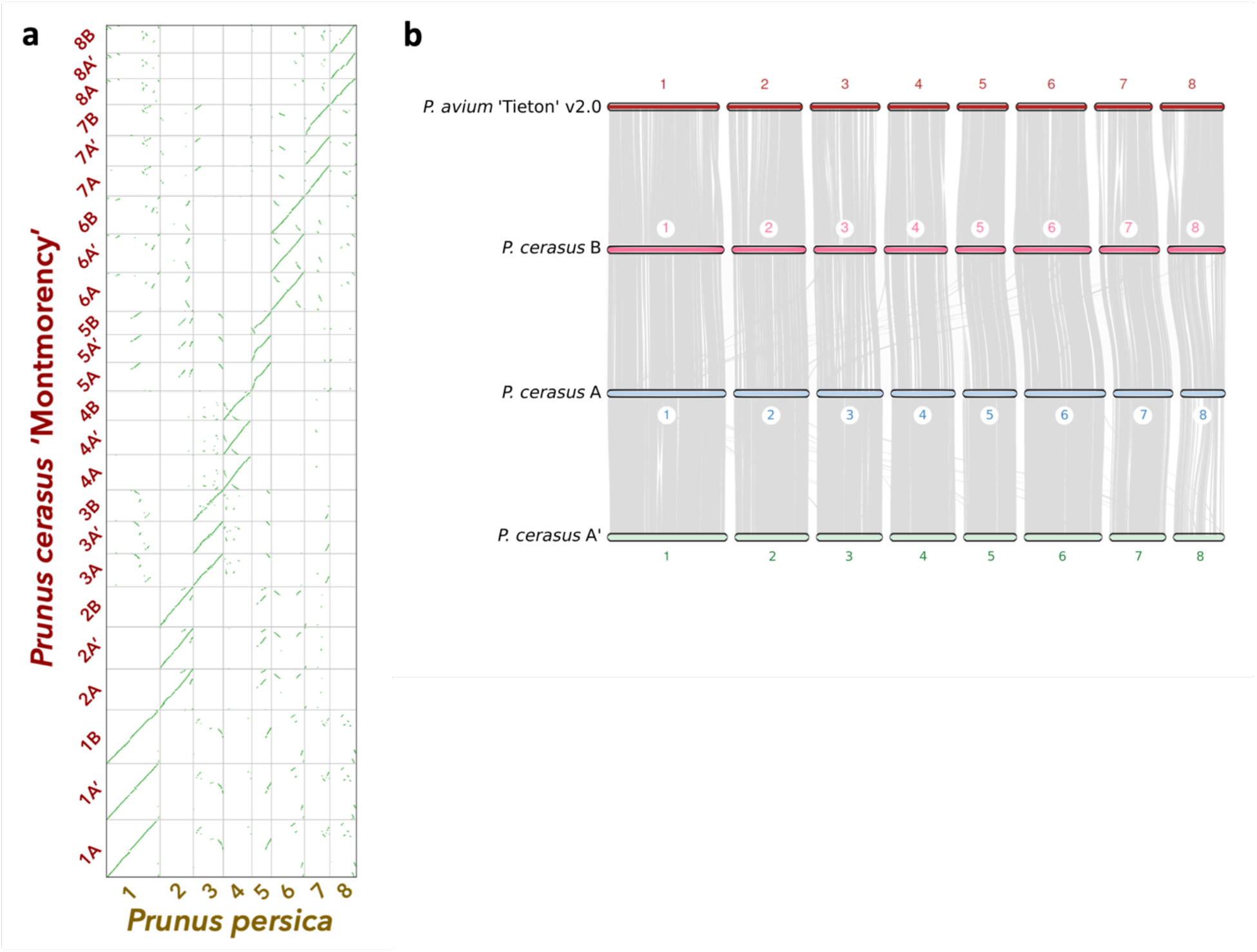
Syntenic relationships between ‘Montmorency’ subgenomes A, A’ and B and other *Prunus* species. a) A syntenic dotplot showing the results of a synteny comparison of the 24 superscaffolds (chromosomes) of *P. cerasus* ‘Montmorency’ and the 8 chromosomes of *P. persica* ‘Lovell’ v2.0 (46). The figure was generated using unmasked coding sequences for each species with the CoGe platform. The prominent linearity for chr #[A, A’, B] for *P. cerasus* vs the respective chr # in *P. persica* highlights the collinearity of this genus and supports the integrity of the assembly. b) Macrosynteny determined with coding sequences shows the three subgenomes of sour cherry are highly syntenic with each other and the published *P. avium* ‘Tieton’ v2.0 genome (22). Each gray line represents a syntenic block between the genomes. Small rearrangements between *P. avium* and each of the Montmorency subgenomes are evident.

**Figure 2:**
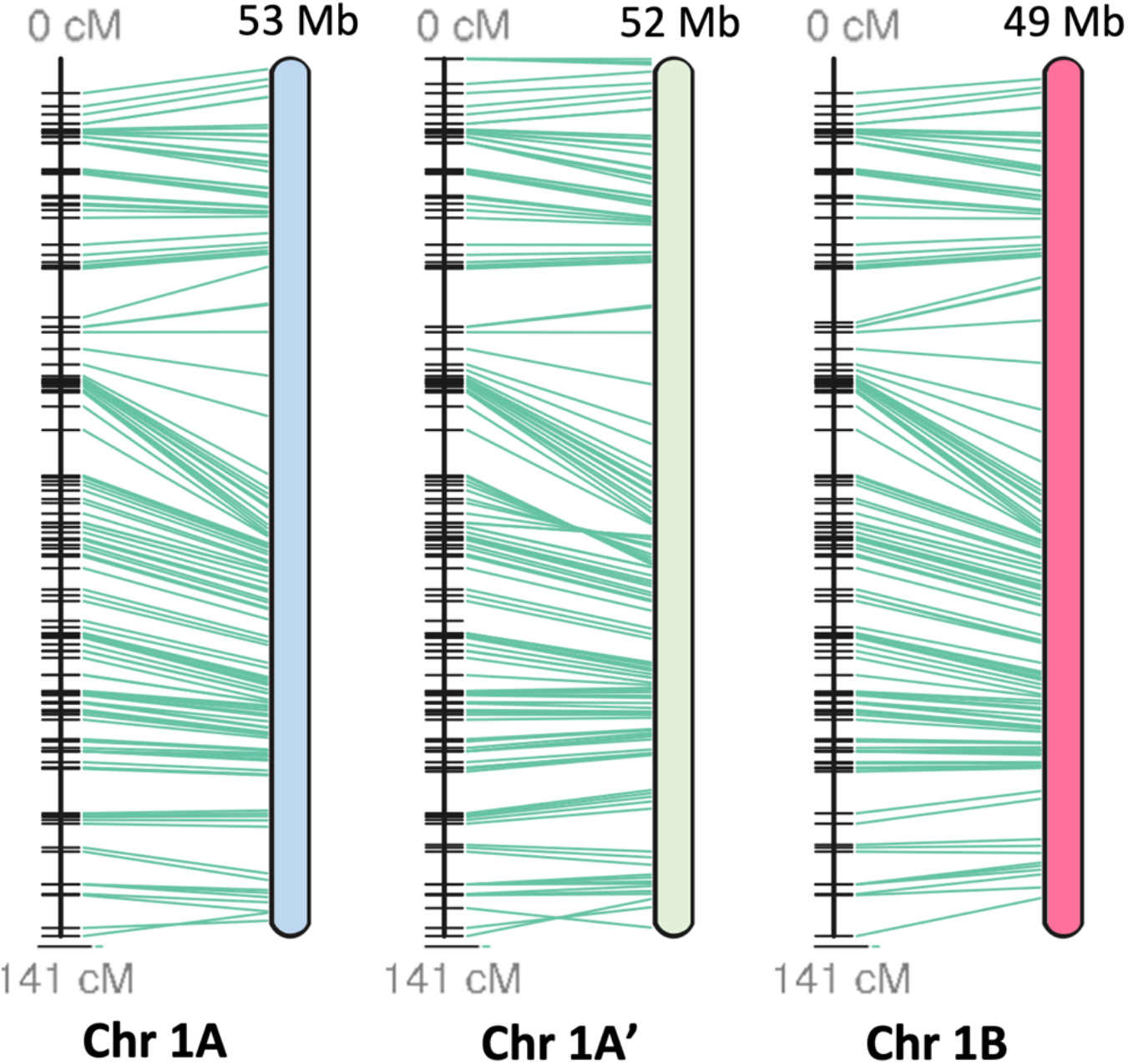
Linearity comparison of linkage group 1 and a published sour cherry genetic map. (11). A total of 545 markers from an F1 sour cherry cross in which ‘Montmorency’ was the female parent were mapped to the assembly and the results demonstrate the high collinearity between the linkage map and assembly. Green lines connect the markers on the genetic map (left) to the physical location in the assembly (right). Each horizontal black line on the genetic map represents one marker. Post-filtering, 426 of the 545 markers mapped exactly once to each subgenome. Subgenome B is a representative of two possible haplotypes. This figure was generated with ALLMAPS (96). Other chromosome sets (2 – 8) are shown in Supplementary Figure 5.

To identify subgenome groups for the 24 chromosomes, we used a strategy based on the rationale that chromosomes enriched for the same set of repetitive elements, represented by k-mer type and abundance, will have a more recent common ancestor. Thus, distinct groups of chromosomes with more k-mers in common may represent subgenomes with the same origin. Unsupervised k-mer clustering to identify allopolyploid subgenomes has been successfully applied to *Miscanthus sinensis* (50), *Nicotiana tobacum* (51), *Triticum aestivum* (51), *Eragrostis tef* (52) and *Panicum virgatum* (53). Using this strategy, we identified 840 25-base pair sequences (25-mers) with more than 10 copies on each chromosome and twice the abundance on any one homoeolog compared to one or both of its sisters. These 25-mers were used to conduct a clustering analysis with all 24 chromosomes of the ‘Montmorency’ assembly, which resulted in two distinct clades: one consisting of 16 chromosomes and the other consisting of the remaining 8 (**Figure 3a, Supplementary Figure 6**). These groupings were mainly attributed to the differential abundance of two 25-mer clusters, designated Group 2 (consisting of 48 25-mers) and Group 3 (consisting of 278 25-mers) (**Supplementary Figure 6**).

**Figure 3:**
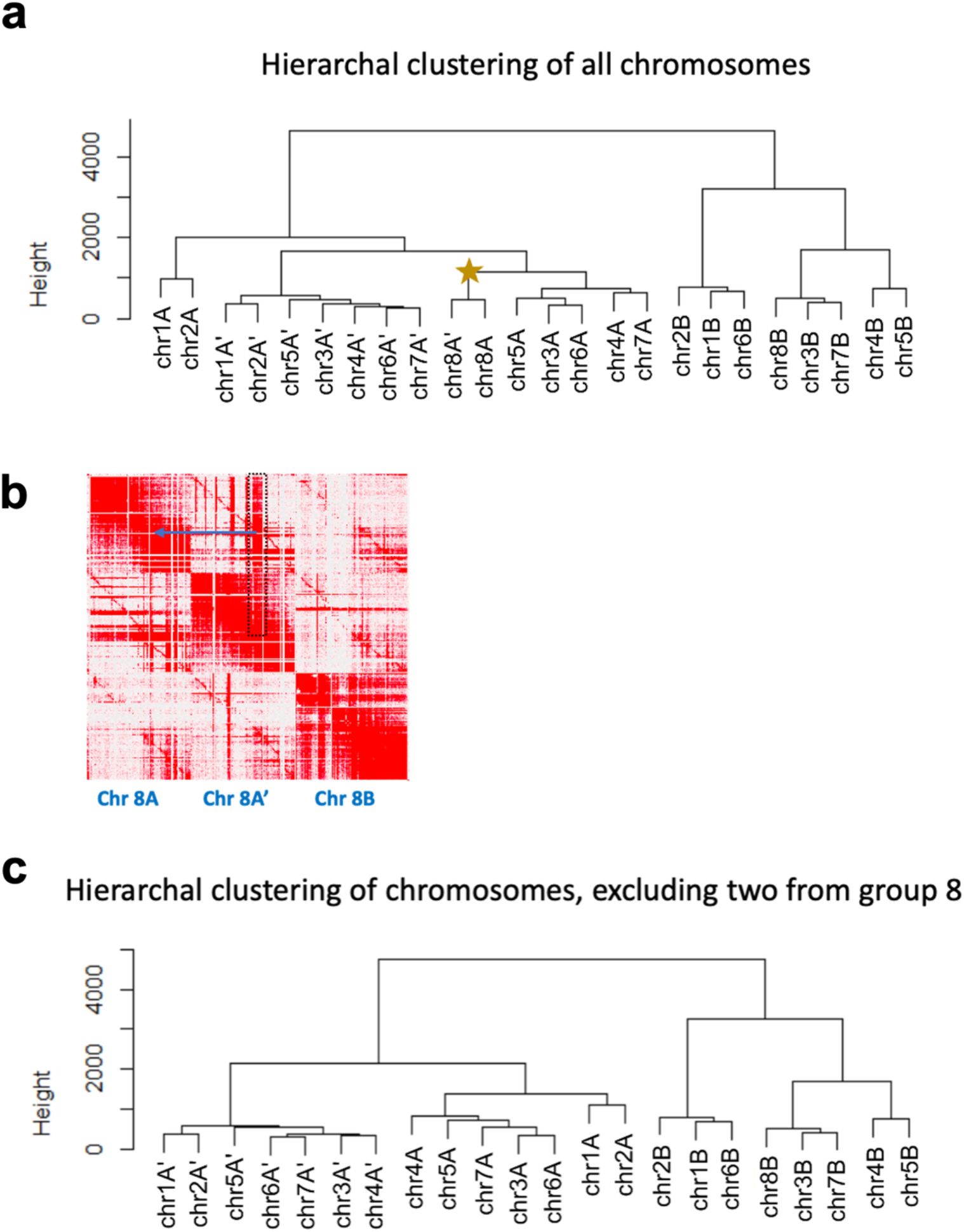
Chr8A and chr8A’ affect the kmer groupings that differentiate subgenomes. a) Hierarchal clustering of all 24 chromosomes based on 25-mer abundance (present 10 times or more on every chromosome and at least twice as abundant on one homoeolog compared to its sisters). The star indicates chr8A and chr8A’. A corresponding heat map is shown in Supplementary Figure 6. b) An enlarged section of the Hi-C matrix for homoeologous chromosome set 8. The dark red signal in the dotted black box indicates an example of sequence on chromosome 8A’ that could have been placed on chromosome 8A in the region indicated with a blue arrow. This could be an assembly artifact (collapse) due to highly similar sequences in these regions, homoeologous exchange between these two chromosomes, or a combination of both. c) Same hierarchal clustering analysis as in a) but excluding the two group 8 chromosomes.

We further posited the set of 16 chromosomes could be subdivided into additional groups given the ease at which most of these chromosomes had been phased. Upon closer examination of the Hi-C matrix of homoeologous group 8, the two included in the group of 16 chromosomes contained many sequences interchangeable with either homoeolog (**Figure 3b**). This may have resulted from the collapsing of similar sequences during assembly, homoeologous exchange, or both. Nonetheless, it was indicative of poorer assembly quality compared to the seven other homoeologous groups and could be clouding the complete separation of these 24 linkage groups into three sets of eight chromosomes. Indeed, when these two linkage groups from homoeologous set 8 were excluded from the clustering, the 22 chromosomes clearly grouped into three clades (**Figure 3c**).

Therefore, a separate clustering analysis including only 14 chromosomes (without homoeologous chromosome 8s) was conducted to better visualize the 25-mers separating the two sets of 7 chromosomes. A total of 481 25-mers in 3 clusters differentiated the two chromosome sets (**Supplementary Figure 7;** groups 5 [n = 44], 6 [n = 148], and 7 [n = 289]). Based on their distinct 25-mer signatures and subgenome assignments (see below), the three chromosome sets were named A, A’ (denoted as A_in all data files), and B. 8A and 8A’ were grouped arbitrarily.

To further investigate the locations of the 25-mer groups contributing to the chromosome subgenome separation, we plotted the densities of several of these 25-mer groups across each chromosome and marked putative centromere locations based on gene and transposable element (TE) densities (**Figure 4a, Supplementary Figure 8a**). Peak 25-mer densities colocalize with the lowest gene content and highest TE density, suggesting these 25-mer groups are most abundant at or near putative centromeres. These estimated centromeric regions agree with a previously-published *P. avium* genome (54). Group 2 and Group 3 densities, which distinguished A/A’ subgenomes from subgenome B in the 24-chromosome clustering analysis (**Figure 3a; Supplementary Figure 6**) clearly exhibited contrasting peak 25-mer densities at approximate centromeres (**Figure 4b, Supplementary Figure 8b**). Likewise, Group 5 and Group 6 25-mers, which differentiated the A and A’ subgenomes (**Supplementary Figure 7**), showed a similar pattern when their densities were aligned to each of the homoeologous chromosome sets (**Figure 4c, Supplementary Figure 8c**). Since highly variable, repetitive satellite DNA is species-specific and often associated with centromere and pericentromeric regions in eukaryotes (55–57), these observations support the claim that A, A’, and B represent distinct subgenomes.

**Figure 4:**
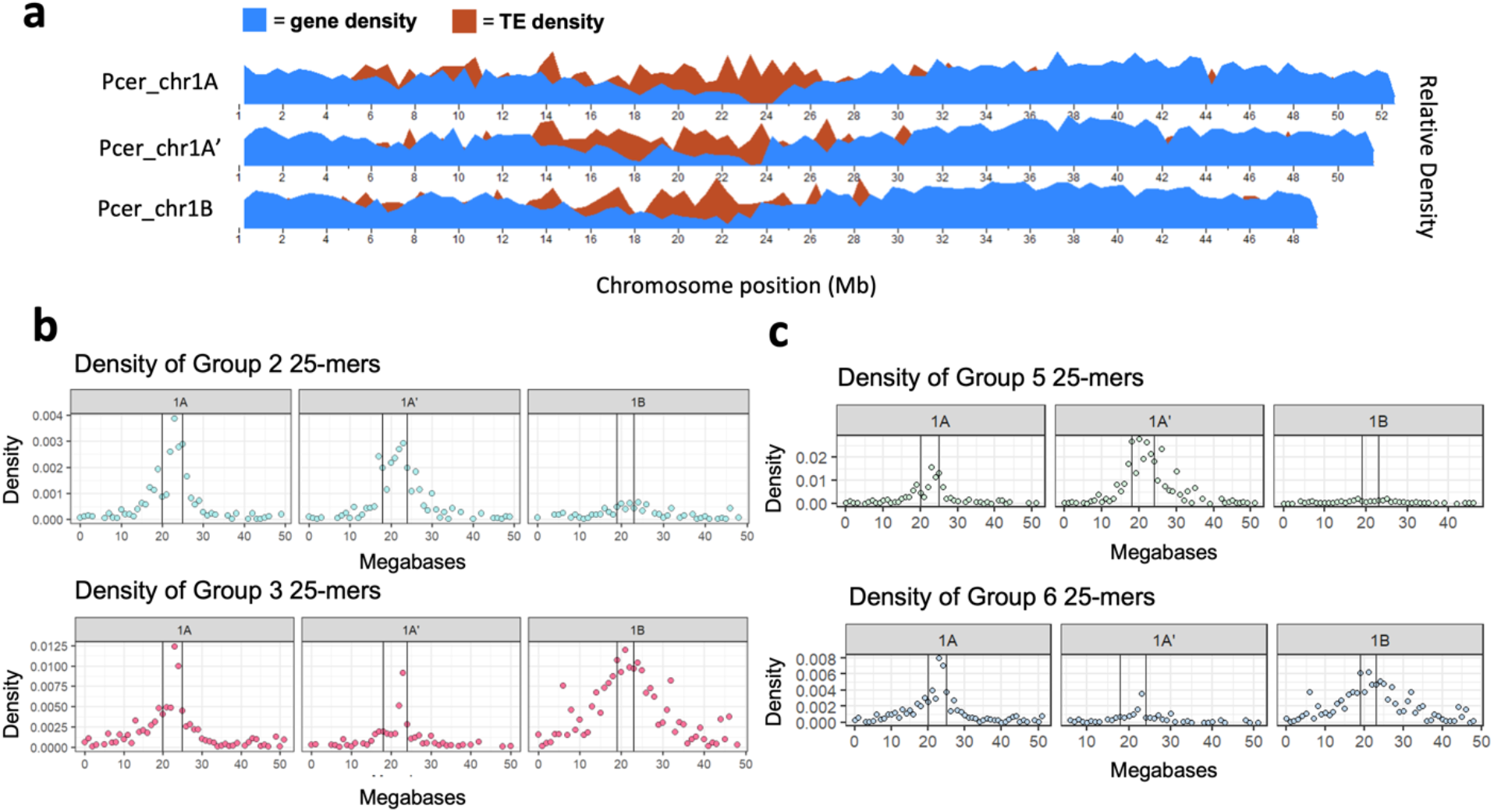
25-mer group densities differentiating the ‘Montmorency’ subgenomes peak at approximate centromeres. Only chr1 is shown for clarity. a) Gene and transposable element (TE) densities plotted along the three chromosome 1 homoeologs. The centromeres are estimated to be regions that coincide with relatively low gene and high TE densities. b) Group 2 and Group 3 25-mer densities (from Supplementary Fig. 6) plotted along the length of chromosome 1. These distinguish the A/A’ subgenomes from subgenome B, when all 24 chromosomes are included in this clustering (Fig. 3a). c) Group 5 and Group 6 25-mer densities (From Supplementary Fig. 7) along the length of chromosome 1. These distinguish A and A’ from one another. In both b) and c), the region between the vertical lines along the density plots designates the approximate location of the centromeres. Corresponding figures for the seven other chromosome sets are in Supplementary Figure 8.

As it is well-established that sour cherry is a tetraploid (5,10,13), we explored the relative dosage of each ‘Montmorency’ subgenome by conducting a depth analysis of the Illumina reads against the assembly. As suspected earlier, this analysis clearly indicated subgenome B had twice the genome dosage of either subgenomes A or A’ as regions along the length of subgenome B frequently showed about twice the read depth of subgenomes A and A’ **(Supplementary Figure 9**). Therefore, the genome structure of *P. cerasus* ‘Montmorency’ is AA’BB. A summary of assembly and scaffolding statistics is given in **Table 1**.

**Table 1:** Summary of the ‘Montmorency’ assembly metrics. Mb = megabases; LG = Linkage Group; NG50 = 50% of the estimated genome size is contained in contigs of equal or greater value; BUSCO = Benchmarking Universal Single-Copy Orthologs (43); LTR = Long Terminal Repeat; TIR = Terminal Inverted Repeat; LAI = LTR Assembly Index (89).

| *P. cerasus* 'Montmorency' assembly metrics |  |  |  |  |  |
| --- | --- | --- | --- | --- | --- |
| **Estimated haploid genome size** (kmer analysis; k = 25) |  |  |  |  | 621 Mb |
| **Estimated Heterozygosity** (total) |  |  |  |  | 4.90% |
| class | aaab | aabb | aabc | abcd |  |
|  | 2.430% | 2.060% | 0.001% | 0.451% |  |
| **Full assembly size** |  |  |  |  | 1066 Mb |
| **Scaffolded assembly size** |  |  |  |  | 771.8 Mb |
| **NG50** |  |  |  |  | 11.56 Mb |
| **Number of contigs** |  |  |  |  | 3592 |
| **Linkage Groups** |  |  |  |  | 24 |
| **BUSCO**<br>**(viridiplantae db10)** | complete | singletons | duplicates | missing | fragmented |
| Scaffolded assembly (24 LGs) | 98.6% | 5.4% | 93.2% | 0.9% | 0.5% |
| subgenome A | 91.5% | 89.6% | 1.9% | 6.4% | 2.1% |
| subgenome A' | 94.8% | 92.9% | 1.9% | 4.0% | 1.2% |
| subgenome B | 90.8% | 88.2% | 2.6% | 7.3% | 1.9% |
| **Estimated % Repeats**<br>**(Full Assembly)** | LTR | TIR | Helitron | Total |  |
|  | 35.6% | 11.6% | 1.3% | 48.5% |  |
| **LAI** | Full assembly (incl. unanchored) |  |  |  | 14.74 |
|  | Scaffolded assembly |  |  |  | 17.09 |

### Annotation of the Prunus cerasus ‘Montmorency’ assembly

For structural annotation of the gene space, RNA-sequencing of a variety of tissues and long-read cDNA datasets were generated for ‘Montmorency’ and used as transcript evidence for de novo gene annotation via MAKER (58,59). MAKER predicted a total of 92,783 protein-coding genes in the full assembly of ‘Montmorency’ after filtering for gene predictions with known Pfam domains (known as the Standard MAKER gene set). The Standard MAKER gene set was the input for the first step of defusion, a MAKER-compatible software designed to disentangle one or more adjacent gene models that are erroneously fused (60). Defusion detected 906 potentially-fused genes in the full assembly; 707 of these were on the scaffolded assembly [chr1A/A’/B – chr8 A/A’/B]. In addition to identifying these candidate fusions automatically with defusion, MAKER was run using only protein evidence to find gene models with more than one protein hit aligning to them – suggesting a possible fusion. Using a custom script, 9867 gene models fitting this criterion were found in the full assembly and 7537 gene models in the scaffolded assembly. Gene models tagged as fusions using these two approaches on the scaffolded assembly were manually checked against protein, RNA, and nanopore cDNA alignments in IGV and added to the breakpoint file if identified as a true fusion (**Supplementary Figure 10**). In total, 4481 genomic regions (gene models) on the scaffolded assembly were locally reannotated using MAKER within defusion. After another Pfam domain search, 9777 of the defused gene models contained known protein domains. These gene models were added back into the final annotation.

Summary statistics for the annotated gene set are given in **Supplementary Table 1**, and **Supplementary Figure 11** shows the cumulative distribution of all gene models’ AED values on the 24 chromosomes of the assembly. Each of the three ‘Montmorency’ subgenomes have similar gene prediction counts and high BUSCO completion scores (85 - 90% for both transcripts and proteins). The shape of AED distribution, with well over half of the gene models having AED values < 0.2, suggests a high-quality annotation that is very agreeable with protein and expressed sequence data.

### Assembly of a Prunus fruticosa draft genome

PacBio long-reads and Illumina short-reads were also generated for a *Prunus fruticosa* accession in the Michigan State University (MSU) germplasm. Similarly to *P. cerasus* ‘Montmorency’, Canu and Pilon were used to create a polished draft assembly. Although the draft assembly is not chromosome-scale, the contigs are large enough for syntenic and gene orthology comparisons (**Supplementary Table 2,** see NG50); and the primary reason for generating this *P. fruticosa* resource was for the assignation of subgenome origins and divergence estimates between progenitors and A, A’, and B in ‘Montmorency’. The *P. fruticosa* draft assembly contains 986 Mb in 3932 contigs, while the genome size was predicted to be 532 Mb. The assembly contains >99% complete BUSCOs, of which 92.7% are duplicated. Additional assembly metrics can be found in **Supplementary Table 2**. A Merqury plot of the *P. fruticosa* assembly revealed common issues associated with polyploid assemblies, namely the collapsing of some haplotypes (red and blue overlap, green and purple overlap) and possible artificial duplication of others (small green overlap with blue and red; **Supplementary Figure 12**). Initially, efforts were made to alleviate these issues and reduce genome complexity by creating a representative haplotype with Purge Haplotigs (44). However, this method was quickly abandoned as it is unknown whether *P. fruticosa* is an allo- or autotetraploid, nor how divergent the alleles are. Given other metrics of the assembly are high and it is prudent to represent the allele diversity for accurate syntelog comparisons, we moved forward to include all contigs in structural annotation. Additional metrics of the draft assembly can be found in **Supplementary Table 2**.

### Annotation of the Prunus fruticosa draft assembly

RNA sequences from five separate tissues from the same *P. fruticosa* accession used for genome assembly, gene models from ‘Montmorency’, and manually-curated protein datasets (61) were used as evidence for annotation of the *P. fruticosa* assembly with MAKER. MAKER predicted a total of 102,361 protein-coding genes for the *P. fruticosa* contigs after filtering to select genes with known Pfam domains and excluding those with known TE domains. The BUSCO completion score for the annotated transcripts of the draft assembly is 97.10% with 92.20% of those duplicated. A summary of statistics for the annotation of the *P. fruticosa* contigs is shown in **Supplementary Table 3.** Like the *P. cerasus* ‘Montmorency’ annotation, well over half of the cumulative fraction of AED values assigned for all gene models (excluding those edited with Apollo), have AED values < 0.2, suggesting most genes are well-supported by transcript and protein homology evidence (**Supplementary Figure 13).**

### Progenitor assignments of the subgenomes in Prunus cerasus ‘Montmorency’

267936 orthologs, or 93.1% of genes in all 7 ‘species’ (*Malus* × *domestica, Prunus persica, P. avium, P. fruticosa*, ‘Montmorency’ subgenome A, subgenome A’, subgenome B) were assigned to orthogroups using OrthoFinder v 2.5.4 (62). Out of all orthogroups, 12051 included at least one ortholog for every species. 336 of these were identified as single-copy orthologous groups. After multiple sequence alignment, trimming, phylogenetic tree construction for every orthogroup, and extraction of progenitor-subgenome gene relationships based on a bootstrap support value >80%, we identified 6797, 7036, and 10398 relationships in subgenome A, A’, and B, respectively. The vast majority of these were syntelogs, and marking their locations colored by representative progenitors on the ‘Montmorency’ chromosomes showed each chromosome was predominantly derived from one progenitor (**Figure 5**). In total, 98.9% (6664/6739) and 99.6% (6957/6984) of syntelog relationships indicated subgenomes A and A’, respectively, were derived from a *P. fruticosa-like* progenitor. Conversely, 98.8% (10245/10370) of syntelog relationships identified for subgenome B supported it as *P. avium*-like. These results also suggested little to no homoeologous recombination had occurred between the A/A’ and B subgenomes.

**Figure 5:**
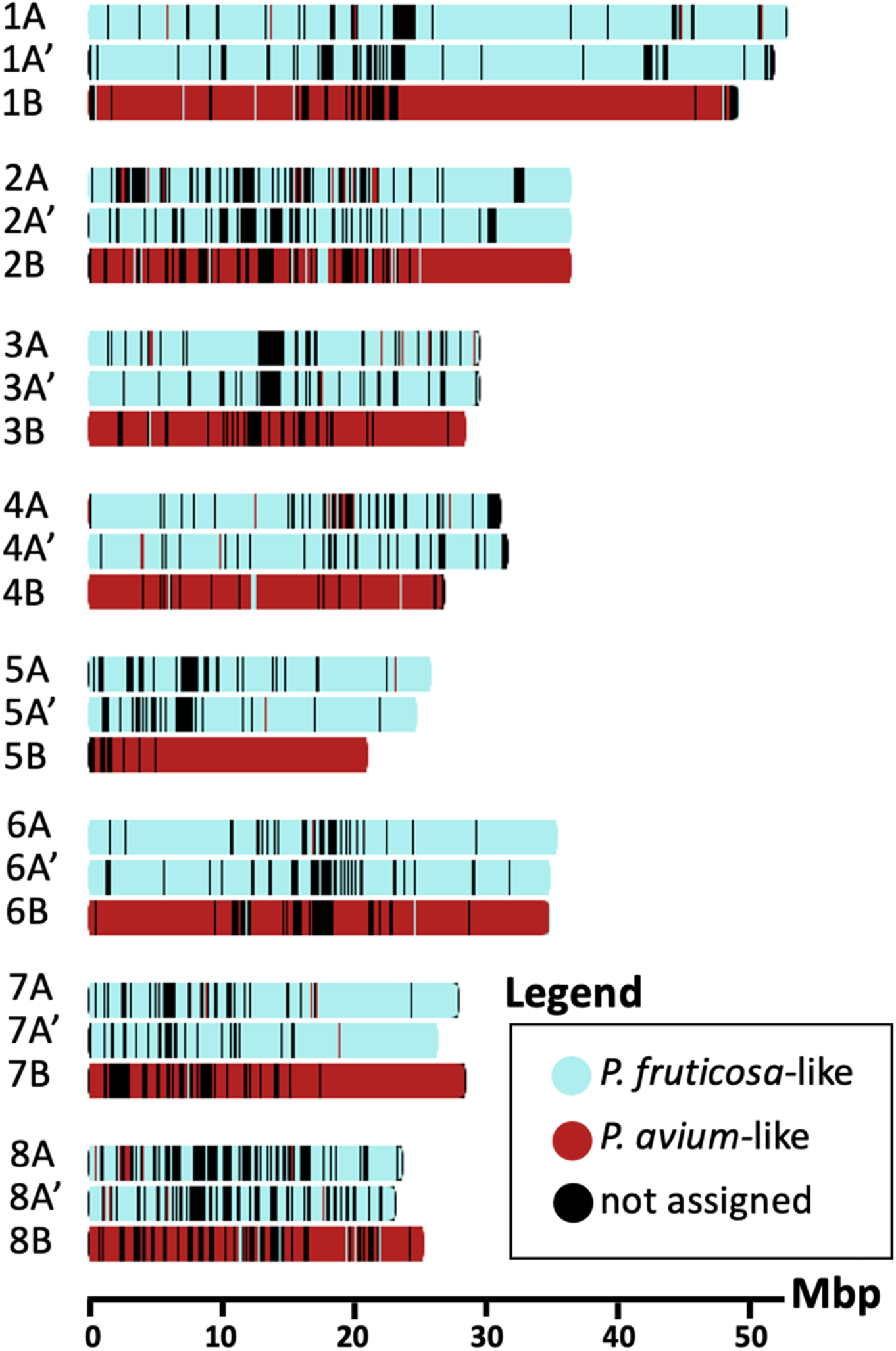
Subgenome assignment using syntelogs reveals little-to-no recombination has occurred between progenitor genomes in ‘Montmorency’. 24,093 syntelogs that were identified with phylogenomic ortholog comparisons and synteny analyses are plotted along the lengths of all 8 chromosome sets and colored by the progenitor they are most likely derived from. Window size for each tick mark ranges between 130 – 133 kilobases and is automatically optimized in chromoMap (reference 133) based on the largest chromosome’s size. Mbp = Megabase pairs.

To identify progenitor relationships of the unanchored genes, we conducted a parallel analysis including the unanchored sequences. However, due to the fragmented nature of these scaffolds, we did not attempt to identify which orthologs were syntelogs. Despite the low number of high-confidence relationships identified in the unanchored sequences (n=858), 87.6% of these genes are *P. avium*-like, further supporting the claim that subgenome B is at twice the dosage of subgenome A and A’.

### DAM gene haplotypes identified in ‘Montmorency’ and Prunus fruticosa

The Dormancy Associated MADS-box genes (DAMs) are six tandemly-arrayed, type II MIKC^c^MADS-box genes required for proper dormancy transitions and bloom time in *Prunus* (26,29). Given the agronomic significance of bloom time, we sought to identify and polish the annotation of these genes in the ‘Montmorency’ and *P. fruticosa* assemblies. BLAST+ analyses (63) revealed DAM gene candidates on ‘Montmorency’ chromosomes 1A, 1A’, and 1B. Upon manual inspection and correction of these gene models using Apollo v 2.6.5 (64), three full haplotypes of DAM1 – DAM6 were found on chr1A, chr1A’, and chr1B. All 18 DAM genes have open reading frames, and all have the characteristic intron-exon structure of these MADS-box genes except for DAM6 on chr1A’. This gene had extremely low expression in the tissues sampled for annotation and only a single, full-length Nanopore cDNA read supported the gene model. It is possible this transcript represents a splice variant, as it is missing 3 of the 9 exons typical of the other DAM genes (29). For the *P. fruticosa* genome, 23 gene models on seven different contigs representing partial and full DAM haplotypes were found; however, only contig 8 and contig 33 contained full DAM haplotypes (i.e., all six DAM genes in tandem), and these regions were confirmed to be syntenic with DAM haplotypes found on ‘Montmorency’ chromosomes 1A, 1A’, and 1B (**Figure 6a**).

**Figure 6:**
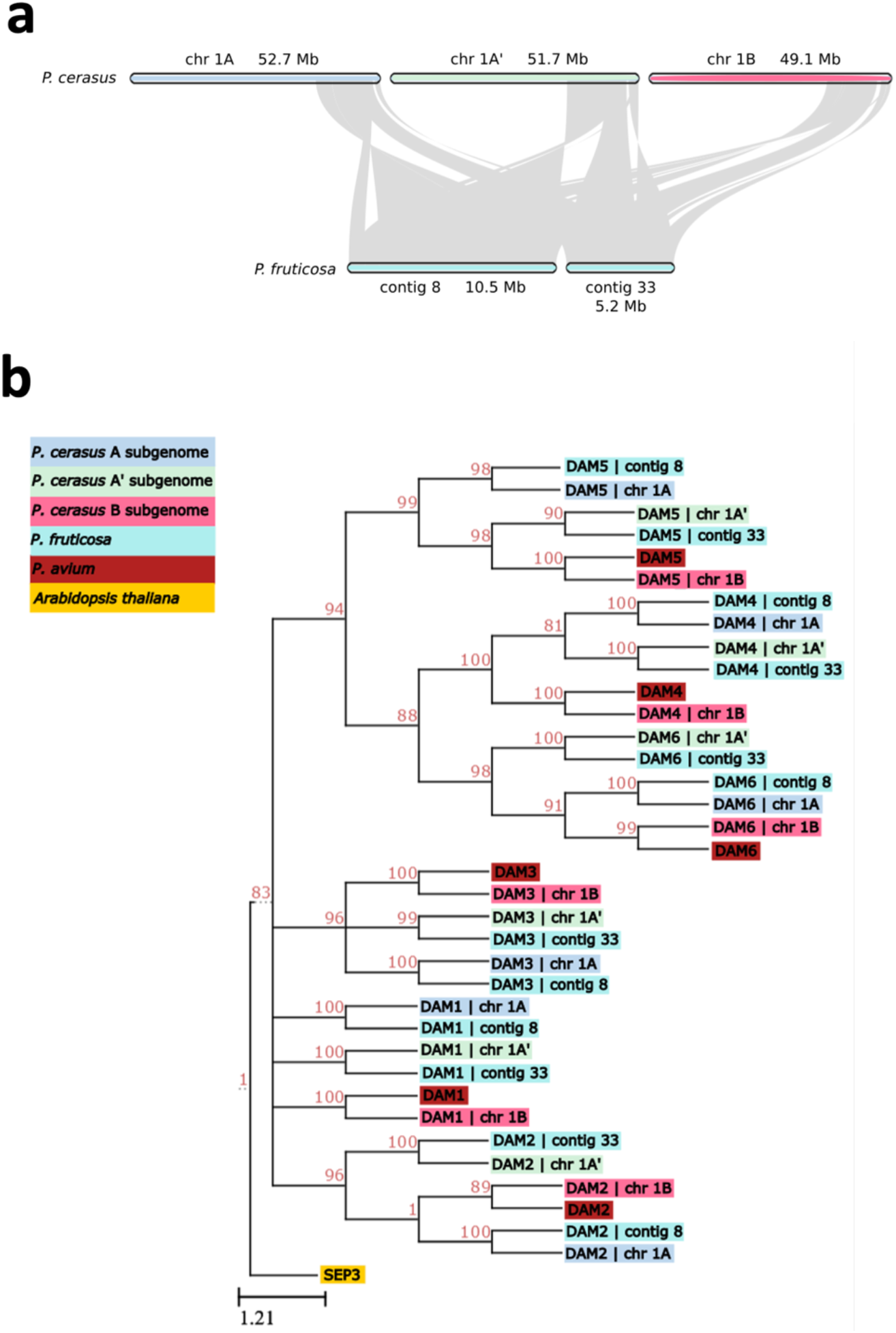
Synteny and phylogeny of the Dormancy Associated MADS-box (DAM) genes in ‘Montmorency’ and *Prunus fruticosa*. a) The genomic regions where full DAM haplotypes were identified in both species show high macrosynteny. b) Phylogeny of the DAM gene coding sequences of ‘Montmorency’, *P. fruticosa*, and *P. avium* (23). *Arabidopsis thaliana* SEP3 was used as an outgroup. The clustering of DAMs together by number suggests correct identification of these genes. The clustering shows the DAM genes from subgenomes A and A’ are most closely-related to *P. fruticosa* DAM genes while DAM genes from subgenome B are most closely-related to *P. avium* DAMs. This agrees with each subgenome’s prior assignment to a *P. avium-like* or *P. fruticosa-like* progenitor. Nodes below an 80% bootstrap value were collapsed.

We next carried out a phylogenetic analysis of all 18 and 12 DAM genes in ‘Montmorency’ and *P. fruticosa*, respectively, with manually annotated DAM genes from *P. avium* (**Figure 6b**) (30). All gene numbers and NCBI identifiers associated with this analysis can be found in **Supplementary Table 4**. All DAM1–DAM6 genes formed well-supported monophyletic clades (BSV >80%) consistent with their order in the tandem array (**Supplementary Figure 14**), garnering further support that these genes were correctly identified. Furthermore, each ‘Montmorency’ DAM gene was sister to the syntelog of its respective progenitor. In other words, all chr1A DAM genes were sister to *P. fruticosa* contig 8 DAM genes, chr1A’ DAM genes were sister to *P. fruticosa* contig 33 DAM genes, and chr1B DAM genes were sister to *P. avium* DAM genes (**Figure 6b**). Interestingly, the gene downstream from the DAM haplotype for ‘Montmorency’ chr1A’ (Pcer_010093) and *P. fruticosa* contig 33 (Pfrut_003731; note that two gene models were created for this region) contain similar insertions of nearly 9 kb in the 6^th^ intron (99.87% identical), providing higher confidence these haplotypes have shared ancestry. The size of this intron on chr1A, chr1B, and *P. fruticosa* contig 8 is approximately 767 bp.

### S-alleles identified in ‘Montmorency’ and Prunus fruticosa assemblies

*S*-alleles, consisting of an RNase (*S*-RNase) and linked F-box protein (SFB), are responsible for gametophytic self-incompatibility in *Prunus* (34). Four *S*-alleles identified in the ‘Montmorency’ assembly on chromosomes 6A, 6A’, and 6B, match the ‘Montmorency’ *S*-alleles previously reported, i.e. *S*_6_, *S*_13m_, *S*_35_, and *S*_36a_ (38,40). The linked SFBs and *S*-RNases comprising these four *S*-allele haplotypes all have >99% identities to their published sequences (37,38,40). BLAST+ results indicated *S*_36a_ is on chr6A, *S*_35_ is on chr6A’, and both *S*_13m_ and *S*_6_ are on chr6B. The two *S* alleles on the same chromosome (6B) is likely an assembly artifact, since these alleles have been demonstrated to segregate independently in sour cherry crosses with ‘Montmorency’ as a parent (40,41). The placement of the *S*_6_ and *S*_13m_ alleles on the *P. avium*-derived subgenome is consistent with the known prevalence of *S*_6_ and *S*_13_ in *Prunus avium* (37). However, the *S*_13m_ in ‘Montmorency’ is a stylar-part mutation of the *P. avium S*_13_ allele that has only been identified in sour cherry (39). The other *S*-alleles in ‘Montmorency’, *S*_35_ and *S*_36a_, have not been identified in *P. avium* and are thought to be derived from the *P. fruticosa*-like progenitor (40). In sour cherry, there are four variants of the *S*_36_ haplotype based on minor sequence differences within and/or flanking the *S*-RNase and SFB coding regions. These four *S*_36_ variants (*S*_36a_, *S*_36b_, *S*_36b2_, and *S*_36b3_) collectively are the most widespread *S*-alleles in sour cherry as all genotypes examined to date have at least one or two *S*_36_ variants (40,65). In the *P. fruticosa* draft assembly, *S*-allele candidates were found on contigs 53 and 1100. Contig 53 was verified to be syntenic to the regions containing the *S*-alleles in ‘Montmorency’ (**Supplementary Figure 15**); however, contig 1100 was too small (130 kb) to do a macrosyntenic analysis using MCScan. Of the sequences used as queries (**Supplementary Table 5**), the closest *S*-RNase and SFB matches for the *P. fruticosa* haplotype on contig 53 were *P. cerasus S*_36b_ variants, with >99% shared sequence identity. All domains characteristic of *S*-RNases were found in the *P. fruticosa S*-RNase identified on contig 53, but a premature stop codon is predicted at amino acid position 67, where a W resides for all other *S*-RNase 36a and b variants (**Supplementary Figure 16a**; note the amino acid fasta sequence reads through the stop codon to indicate conserved domains). The SFB protein on contig 53 has an open reading frame, expected protein domains, and an amino acid sequence identical to the *P. cerasus* SFB_36b_ sequence (**Supplementary Figure 16b**) (40). In summary, *S*_36_ variants have not been previously identified in *P. fruticosa*, so the *S*_36_ variant found in this *P. fruticosa* draft assembly supports the hypothesis that sour cherries, including ‘Montmorency’, inherited their *S*_36_ haplotype(s) from a *P. fruticosa-like* progenitor.

### Prunus cerasus ‘Montmorency’ is descended from a recent hybridization event

Lastly, we sought to estimate divergence time of each ‘Montmorency’ subgenome from its most closely related representative progenitor. First, we examined the individual topologies of the 336 phylogenetic trees of single-copy orthologs. Only single-copy orthologs were used in these analyses to diminish the effect recent gene duplication events would have on divergence estimates. Based on previous phylogenetic assessments (1–3,66) and the ‘Montmorency’ subgenome assignments, a topology with A or A’ sister to the *P. fruticosa* ortholog, B sister to *P. avium, P. persica* sister to the cherries, and *Malus* × *domestica* sister to *Prunus* would most accurately estimate when each subgenome last shared a common ancestor with its representative progenitor (i.e., when these lineages began diverging). The hybridization event ‘Montmorency’ is descended from would have occurred sometime after this estimate.

Because *P. fruticosa* is a verified tetraploid, the ortholog in each topology is an assembly collapse of four possible alleles. If *P. fruticosa* is an allotetraploid with two intact and divergent subgenomes, approximately half of the single-copy orthologs would be expected to come from the first subgenome while the other half would come from the second subgenome. If A and A’ share a more recent common ancestor with one *P. fruticosa* subgenome compared to the other, we would expect the orthologs from A and A’ to be sister to *P. fruticosa* in an approximately equal number of trees. Moreover, if A and A’ are descended from divergent *Prunus* ancestors, we would rarely expect their single-copy orthologs to be sister to one another.

The most frequent topologies observed among the single-copy orthogroups include one where A is sister to *P. fruticosa* (n=43) and the other where A’ is sister to *P. fruticosa* (n=70; **Supplementary Figure 17**). In all single-copy ortholog topologies, A was sister to the *P. fruticosa* ortholog 42% of the time (n=142), whereas A’ was sister to the *P. fruticosa* ortholog 49% of the time (n=165). A was sister to A’ in only 1.8% of topologies (n=6). These results are consistent with *P. fruticosa* being an allotetraploid, and that A and A’ are diverged enough to be considered derived from separate *Prunus* species. Subgenome B was sister to *P. avium* in nearly all single-copy ortholog trees (n=310; 92%).

The r8s analysis of the most frequent topologies suggested ‘Montmorency’ subgenome A and *P. fruticosa* began diverging 1.61 – 1.63 mya (topology B, node 6), subgenome A’ and *P. fruticosa* began diverging 4.48 – 4.51 mya (topology A, node 4), and subgenome B and *P. avium* began diverging < 1.72 mya (topology A, node 6; topology B, node 5; **Supplementary Figure 17**). Taken together, these results suggest the hybridization event *P. cerasus* ‘Montmorency’ is descended from occurred less than 1.61 mya.

## Discussion

Here we report the genome assembly for sour cherry (*Prunus cerasus*) cultivar Montmorency, which to our knowledge is the first genome sequence for sour cherry to be published. We chose to sequence ‘Montmorency’ as it is the most commonly grown cultivar in the United States. This chromosome-scale assembly is highly collinear with a published sour cherry genetic map (11), is syntenic with other *Prunus* species (22,46), and has a quality annotation with thorough manual curation. Additionally, we assembled and annotated a draft genome of *P. fruticosa*, the closest extant relative of one of sour cherry’s proposed progenitor species. We expect both resources to be informative for future sour cherry breeding strategies and comparative genomic studies in *Prunus*.

The allotetraploid origin of sour cherry is well-established and further supported by this work; however, the three subgenome composition of ‘Montmorency’, AA’BB, with two divergent genomes contributed by a *P. fruticosa*-like ancestor, was an unexpected result. The separation of subgenomes A and A’ using k-mers required the exclusion of chromosomes 8A and 8A’, which showed lower assembly quality compared to other chromosomes. In addition to k-mer evidence, the low frequency in which A and A’ genes were sister to one another in single-copy ortholog trees lends strong support for treating them as separate subgenomes derived from different *Prunus* species. Until now, concrete evidence of the origin of *P. fruticosa* has been lacking despite the recent publication of its genome sequence (17). Our results suggest *P*. *fruticosa* is an allopolyploid hybrid of two distinct *Prunus* species. However, since large-scale evolutionary studies of *Prunus* rarely include this species (1–3), the extant relatives of its progenitor species are unknown and is a question for future research. Identification of the *P. fruticosa* progenitor species (or, more probably, their extant relatives) would allow for greater resolution of the dynamics of subgenome A and A’ within sour cherry.

The subgenome assignments for ‘Montmorency’ were further supported by our results for two sets of biologically significant genes for *Prunus* in the ‘Montmorency’ reference and *P. fruticosa* draft genomes: the DAM genes and *S*-alleles. Alleles of both gene sets were consistent with the conclusion that ‘Montmorency’ has an AA’BB subgenome structure where A/A’ and B are derived from *P. fruticosa*-like and *P. avium*-like ancestors, respectively. In this work, we document the first discovery of an *S*_36_ variant in *P. fruticosa*, supporting the previous hypothesis that *S*36 variants identified in sour cherry are derived from a *P. fruticosa-like* progenitor (40). An interesting question moving forward is how the progenitor origins of the DAM genes may relate to bloom time in diverse sour cherry accessions. Sour cherry exhibits a transgressive range in bloom time: some genotypes bloom earlier or later than either progenitor (11). *P. fruticosa* is native to colder northern latitudes in eastern Europe and flowers later than *P. avium*, which originates from the Mediterranean region south of the Black Sea (67). One might expect to see selective pressure on the DAM genes depending on whether a *P. cerasus* genotype distributes to more northern or southern latitudes, tailoring bloom time to maximize reproductive potential. However, this question remains largely unexplored.

Our subgenome assignments for ‘Montmorency’ also provide insights into the possible gametes that formed ‘Montmorency’. First, the ‘Montmorency’ subgenomes derived from the *P. fruticosa-like* progenitor (A/A’) do not show any evidence of recombination with the subgenome derived from the *P. avium*-like progenitor (B). This lack of recombination supports the theory that ‘Montmorency’ was formed by one gamete from the *P. fruticosa*-like ancestor and one gamete from the *P. avium*-like ancestor. Second, since A and A’ were readily distinguishable according to gene and repetitive sequence differences (with the exclusion of chr8A and chr8A’), this implies very few homoeologous exchanges (mixing of genomes) have occurred between these more similar subgenomes. This finding once again suggests the *P. fruticosa*-like progenitor was an allopolyploid and the gamete that gave rise to ‘Montmorency’ resulted from preferential chromosome pairing of the two subgenomes. Chr8A and 8A’ may be an exception; however, it is unclear whether this is due to subgenome mixing, disproportionately high sequence similarity among these chromosomes, assembly artifacts, or a combination of one or more scenarios. These findings are consistent with the possibility that ‘Montmorency’ was formed from the fusion of a reduced gamete from a *P. fruticosa*-like ancestor and an unreduced gamete from a *P. avium*-like ancestor (8). In the case of ‘Montmorency’, chloroplast data identified the *P. fruticosa-like* progenitor as the maternal parent (6).

It follows that the integrity of the three ‘Montmorency’ subgenomes is not likely to be transmitted to the next generation. If the A and A’ chromosomes preferentially pair, crossing over will result in a patchwork of regions exchanged between these two homoeologous chromosome sets. Additionally, cytological analysis of meiotic pairing for ‘Montmorency’ shows lack of complete bivalent pairing: quadrivalents, trivalents and univalents occur (10,68). Segregation data support primarily subgenome pairing (A with A’ and B with B); however genetic results are consistent with occasional homoeologous pairing (5,11). This suggests that homoeologous exchanges could occur between the A and A’ subgenome chromosomes and the B subgenome chromosomes, further eroding the subgenome integrity in ‘Montmorency’ offspring.

Poor fertility supported by evidence of irregular meiosis and occasional tetrasomic inheritance are prevalent in sour cherry germplasm (69). In general, the quantity of fruit set is vastly below what the plant could support. Commercial cultivars such as ‘Montmorency’ are the exception as this cultivar can achieve a “full crop” by setting approximately 30% of its fruit. Indeed, it is the high fertility in ‘Montmorency’ that led it to be the predominant cultivar in the United States. However, even in crosses between two productive cultivars, the low fertility in the offspring is clear. For example, in one study (9), the mean fruit set and pollen germination for the German sour cherry cultivar Schattenmorelle and the Hungarian cultivar Érdi Bötermö was 16.0% and 13.4%, respectively for fruit set, and 18.5% and 8.0% respectively for pollen germination. When these two cultivars were crossed, the mean values for fruit set and pollen germination of the 86 offspring were just 6.8% and 6.6%, respectively. Both these values are far below what is needed for a commercial crop. As such, breeding for higher fruit set is a major but challenging goal for the MSU sour cherry breeding program.

In the present study, we provide evidence that the allotetraploid event from which the ‘Montmorency’ lineage is descended occurred less than 1.61 million years ago. Though not considered recent by some standards, sour cherry is a long-lived perennial species; therefore, while 1.61 million years represents many generations and opportunities for recombination for an annual species, the same time span equates to far fewer generations for sour cherry. For example, the first written record of ‘Montmorency’ was in the 17th century, making it ~ 400 years old (70). Thus, this sour cherry lineage may be considered younger than short-lived polyploids in terms of absolute number of generations. It must be noted, however, that the progenitor representatives used in dating analyses may greatly affect estimates. Additionally, since sour cherry is a species thought to have formed multiple times (6), the timing of hybridization for certain lineages would be expected to vary. Still, given these results and speculations, it would be reasonable to suggest sour cherry exemplifies the behavior of a neopolyploid actively undergoing the process of diploidization. Cytological and genetic data is consistent with a neopolyploid as irregular meiosis visualized as trivalents and quadrivalents is documented in several sour cherry genotypes (10,68), and genetic data indicates that although disomic inheritance is more common, tetrasomic inheritance has also been observed (9,11,71). Such events can result in aneuploid gametes, endosperm imbalance, embryo abortion, and unsuccessful seed and fruit development. Neoallopolyploids are especially prone to such issues depending on the likeness of their progenitors’ genomes (72). If progenitors are relatively divergent, preferential pairing of homologous chromosomes and *not* homoeologous chromosomes will more likely occur, and the polyploid will exhibit frequent bivalent formation during meiosis and diploid-like segregation.

However, if progenitors are more similar in terms of their sequence divergence and collinearity, homoeologous chromosomes may exchange genetic information and create imbalanced genomic combinations. This could result in pairing irregularities during meiosis and reduced fertility in subsequent generations.

Theoretically, over time, selection would act swiftly against these infertile genetic combinations and the polyploid would eventually ‘diploidize’ (72–74). However, the diploidization process in sour cherry has likely been repressed by the prevalent intercrossing with both its progenitor species, as natural hybrids among the three cherry species (*P. cerasus*, *P. avium*, and *P. fruticosa*) are common (13). In certain scenarios, progeny from a cross of sour cherry (AA’BB) and *P. avium* (BB) or *P. fruticosa* (AAA’A’) would likely inherit imbalanced subgenome combinations. Furthermore, human selection may also have constrained the diploidization process. For example, in Hungary and Romania, human selection of vegetatively-propagated century-old landrace cultivars Pándy and Crişana, respectively, favored fruit quality over fruit quantity, as these landrace cultivars have extremely low fruit set but highly desirable fruit quality (75). Taken together, the evolutionary history of sour cherry illustrates the intersection of fundamental principles of natural selection and human influence.

Finally, an intriguing question moving forward is how many separate allotetraploid lineages led to what we consider to be sour cherry. Chloroplast data indicates sour cherry was independently formed at least twice: although *P. fruticosa* is more commonly the maternal parent, cultivars with *P. avium* as the maternal parent have also been identified (6). As the native distributions of *P. fruticosa* and *P. avium* overlap, sour cherry could have been formed by a reduced gamete from *P. fruticosa* and an unreduced gamete from *P. avium*, as speculated above for ‘Montmorency’. However, it is also possible sour cherry lineages could be the product of a triploid bridge. In this scenario, a hybrid of a *P. fruticosa-like* and *P. avium-like* species (*Prunus* × *mohacsyana*) would produce an unreduced gamete (3x) that either fertilized or was fertilized by a *P. avium* gamete (1x) to form allotetraploid *P. cerasus*. Irrespective of these different evolutionary trajectories, it is important to consider the ‘Montmorency’ subgenome structure reported herein, may be unique to this genotype and therefore may not represent the subgenome structure of a range of sour cherry accessions.

## Conclusions

We present the ‘Montmorency’ reference genome and a *P. fruticosa* draft genome to be used for future comparative studies in the genus *Prunus* and beyond. Our characterization of the ‘Montmorency’ subgenomes provides a valuable resource for exploring the evolutionary history of sour cherry with wider implications for questions surrounding allopolyploidization and neopolyploidy. These resources will aid in developing targeted breeding strategies for sour cherry and allow investigation into whether an imbalanced subgenome composition leads to the low fruit set prevalent in the species.

## Methods

### Collection of materials

Young leaves for gDNA libraries were collected fresh (for Hi-C, *P. cerasus* ‘Montmorency’ only) or flash-frozen in liquid nitrogen and stored at −80°C until extraction (for both PacBio SMRT sequencing and Illumina HiSeq) from a clone of ‘Montmorency’ and an accession of *P. fruticosa* growing at Michigan State University’s Clarksville Research Center in Clarksville, Mich., in the spring of 2019. Tissues for RNAseq and Nanopore cDNA libraries were collected the same year from ‘Montmorency’ and flash frozen in liquid nitrogen and stored at −80°C until extraction. For *P. cerasus* ‘Montmorency,’ biological replicates were collected for RNA extraction and RNAseq / Nanopore cDNA-sequencing from the tissues indicated in **Supplementary Figure 18**. Fruit tissue was collected and staged based on the double-sigmoidal growth curve characteristic of *Prunus* sp. (76,77). Vegetative and floral meristems in typical floral positions were broadly characterized as pre-floral-initiation, transitioning to floral, and organ differentiation according to histological sectioning (data not shown). For *P. fruticosa*,biological replicates were collected for extraction and RNAseq from young leaves, whole flowers at balloon stage, whole fruits in stage I, whole fruits at the end of stage II, and whole fruits in stage III.

### DNA extraction, library preparation, and sequencing

Extraction of high molecular weight (HMW) DNA from young leaves was done at the University of Georgia’s Genomics and Bioinformatics Core using a nuclei-extraction method for both *P. cerasus* ‘Montmorency’ and *P. fruticosa*. From this HMW DNA, a large SMRTbell library (>30kb) was prepared and sequenced on six flow cells of a PacBio Sequel II machine for each species, producing 61.98 Gb of data (100X coverage of 621 Mb estimated genome size) for *P. cerasus* ‘Montmorency’ and 48.13 Gb of data (90.5X of 532 Mb estimated genome size) for *P. fruticosa*. A second batch of young leaves was used to extract DNA for short-read sequencing using the DNEasy Plant kit (Qiagen, Valencia, CA, USA). An Illumina TruSeq gDNA library was prepared and sequenced for both species on a HiSeq4000 at the Research and Technology Facility (RTSF) of Michigan State University (MSU), and approximately 34.7 Gbp of data was produced (56X coverage) for *P. cerasus* ‘Montmorency’ and 40 Gbp of data was produced for *P. fruticosa* (75.2X coverage). A third collection of fresh young leaves from *P. cerasus* ‘Montmorency’ was shipped overnight on ice to Phase Genomics (Seattle, WA), where a Hi-C Proximo Library was created with a DpnII restriction enzyme. The Hi-C library was 150 bp paired-end sequenced on a HiSeq4000 instrument at MSU’s RTSF, producing 93.3 Gbp of data or 150.5X physical coverage.

### RNA extraction, library preparation, and sequencing

All RNA was extracted using a CTAB-based protocol (78). One Illumina TruSeq Stranded mRNA library was prepared for each of the 2 – 3 biological replicates per tissue used for RNAseq (**Supplementary Figure 18**, excl. c, d, g, j-l for ‘Montmorency’). Young leaves, whole flowers at balloon stage, whole fruits in stage I, whole fruits at the end of stage II, and whole fruits in stage III, or RNA from 5 tissue-types total, were sequenced for *P. fruticosa* at MSU’s RTSF. 24 RNAseq libraries for ‘Montmorency’ and 14 RNAseq libraries for *P. fruticosa* were 150 bp paired-end sequenced on a HiSeq4000, producing between 25 and 36.6 million reads per library. One library from ‘Montmorency’ (replicate of whole fruits at the end of stage II) and one library from *P. fruticosa* (replicate of whole fruits at stage I) were deemed contaminants / low-quality based on unusually poor alignment to the respective assemblies and were excluded from downstream analyses. In addition to most of the tissue types used for RNAseq (excluding mature fruit mesocarp and mature fruit exocarp), whole fruits at the beginning of phase II, vegetative apices in early summer, transitioning apices in midsummer, mature leaves in late spring, mature leaves in midsummer, and floral apices during organ development (confirmed via histological observations – unpublished results) were included in the Nanopore cDNA-sequencing (12 tissue-types total) for ‘Montmorency’ (**Supplementary Figure 18** c, d, g, j-l).

### Assembling the genome of Prunus cerasus ‘Montmorency’

Illumina gDNA reads’ 25-mers were counted using Jellyfish v 2.2.10 (RRID:SCR_005491) (79) and the resulting histogram was visualized using GenomeScope v2.0 (RRID:SCR_017014) (80). Canu v 1.9 (RRID:SCR_015880) (81) was used to assemble the PacBio reads. Reads below 5kb in length were excluded from the assembly process and batoptions were set to `“-dg 3 -db 3 -dr 1 - ca 500 -cp 50”` to utilize the heterozygosity to assemble all possible haplotypes. The assembly was polished with the Illumina gDNA reads iteratively 4 times with Pilon v1.23 (RRID:SCR_014731) (82). Reads were aligned to the assembly with Bowtie2 v 2.3.4.2 (RRID:SCR_016368) (83) until there was no further improvement in read alignment. The polished assembly was visualized with Bandage (RRID:SCR_022772) (84), a BUSCO analysis was done to assess gene space (RRID:SCR_015008) (43), and Merqury (RRID:SCR_022964) (42) was used to determine phasing quality and genome-completeness.

### Scaffolding the Prunus cerasus ‘Montmorency’ assembly

Preliminary scaffolding results indicated two full haplotypes (16 pseudomolecules) had been well-assembled while a third (8 pseudomolecules) experienced sudden and frequent drops in Hi-C signal along the diagonal, likely due to haplotype switching and the 3D-DNA software attempting to position multiple, non-collapsed alleles next to collapsed sequence (i.e., heterozygous bubbles). The rest of the assembly was considered unanchored. We posited that removal of similar haplotypes would lead to a tidier representation of the third group. Purge_haplotigs v1.1.2 (44) was used with a cutoff value of 99% alignment to set aside very similar alleles in the assembly prior to scaffolding. Since the contig lengths of the 16 well-assembled pseudomolecules were much larger than the other 8, the lengths of the purged contigs were manually inspected to avoid removal of this sequence. As a result, purged contigs greater than 400kb were added back into the assembly prior to scaffolding. After this size selection, we verified we had removed mostly alternative alleles of the three assembled groups in two ways. First, BUSCO analyses showed completeness to be very high (>90%) and duplication to be very low (<3.0%) for each of the three pseudomolecule groups (**Table 1**). Second, a k-mer assessment using Merqury indicated 25-mers in the remaining purged contigs were found only once and at all relative multiplicities (1X - 4X) in the Illumina read dataset. If 25-mers at any multiplicity had been present in the purged contigs more than once, this would suggest more than one haplotype had been removed from the assembly (**Supplementary Figure 3**). After reducing the complexity of the assembly as described above, Hi-C reads were aligned to the assembly using BWA v 0.0.7.17 (RRID:SCR_010910) (85) within the Juicer pipeline (RRID:SCR_017226) (86). The -S flag was set to exit the pipeline early after production of the merge_nodups.txt file. This file was then used as input for 3D-DNA (RRID:SCR_017227) (87), which was run with `“--editor-repeat-coverage 5”` to prevent areas of the assembly with higher levels of coverage (due to ploidy) being flagged as “junk.” The output was a Hi-C matrix (.hic) that was manually edited in JuiceBox Assembly Tools (RRID:SCR_021172) (88) to correct misassemblies. Following manual editing, the new .hic and .asm files were used to create the chromosome-level .fasta file with the script run-asm-pipeline-post-review.sh from the 3D-DNA suite of tools. The resulting superscaffolds (chromosomes) were named according to a syntenic comparison with a peach genome (45,46) and subgenome assignments based on 25-mer groups.

### Assessing repeat content and quality

The LTR assembly index (LAI) (89) was determined by first identifying TEs with LTR_FINDER_parallel (RRID:SCR_018969) (90) and LTRharvest (91), then combining the output files and using it as input for LTR_retriever (92). The EDTA pipeline was used to estimate repeat content and produce a custom repeat library (RRID:SCR_022063) (93).

### Marker mapping and visualization

582 marker sequences from a genetic map of an F1 cross of two sour cherries (‘Montmorency’ × 25-02-29, n = 53) (11) were downloaded from the Genomic Database for Rosaceae (GDR; https://www.rosaceae.org/) (94,95) and mapped to the 24 superscaffolds of the assembly using BLAST+ v 2.2.31(RRID:SCR_011820) (63). Markers mapping more than four times or below 80% of their length were filtered from the dataset, resulting in 545 unique markers’ mappings visualized in ALLMAPS (RRID:SCR_021171) (96).

### Annotation of the genome of Prunus cerasus ‘Montmorency’

We used multiple sources of high-quality data to annotate the *P. cerasus* ‘Montmorency’ genome, including RNAseq and long-read cDNA-PCR sequencing using a GridION machine (Oxford Nanopore Technologies), and manually curated protein databases (61,97). All data were processed to produce .gff3 files which were used as input for MAKER (RRID:SCR_005309) (59).

### Preparation of RNAseq data for MAKER

Adapters and low-quality bases were removed from RNAseq reads (2-3 reps per tissue, 23 libraries total) with Trimmomatic v 0.39 (RRID:SCR_011848) (98). Individual libraries, totaling 3.1 billion reads, were aligned to the ‘Montmorency’ genome assembly using default parameters in STAR v 2.7.3a (RRID:SCR_004463) (99). Alignment rates were 94%+ per library. Approximately 33-39% of reads mapped to multiple locations, and random checks of several alignments confirmed these reads were aligning to homoeologous chromosomes and/or alleles (e.g., 1A and/or 1A’ and/or 1B). SAMtools v1.9 was used to sort and index all .sam/.bam files (RRID:SCR_002105) (100). All alignments were merged, and a transcriptome assembly was created using StringTie v 2.1.2 (RRID:SCR_016323) (101). The transcriptome assembly was checked against raw RNAseq and protein alignments for improper fusions and breaks in potential genes with the Integrative Genomics Viewer (IGV) v 2.8.0 (RRID:SCR_011793) (102), and parameters were adjusted accordingly in StringTie v 2.1.2 (`“-m 200 -t -c 3 -f 0.05 -g 50”`).

The final .gtf file was converted to .gff3 using the gffread function in Cufflinks v 2.2.1 (RRID:SCR_014597) (103) and the features ‘StringTie,’ ‘transcript,’ and ‘exon,’ were replaced with ‘est2genome,’ ‘expressed_sequence_match,’ and ‘match_part,’ respectively, for compatibility with MAKER v 2.31.10 (59).

### Preparation of long-read RNA sequencing for MAKER

Nanopore reads were demultiplexed, trimmed, and filtered (reads <150bp were dropped) with Porechop v 0.2.4 (RRID:SCR_016967) (104) and NanoPack (105). 4.8 million reads were aligned at a rate of 89% to the ‘Montmorency’ genome assembly using minimap2 v 2.15 (RRID:SCR_018550) (106) with the following parameters: `“-N 5 -ax splice -g2000 -G10k”`. Sorting and indexing of .sam and .bam files was done with SAMtools v. 1.9. The transcriptome assembly was built using StringTie2 (`“-m 150 -t -c 1 -f 0.05 -g 50”`), and the .gtf was converted to .gff3 and features changed similarly to the RNAseq data prior to giving the data to MAKER.

### Preparation of protein data for MAKER

Manually curated Uniprot viridiplantae protein sequences (RRID:SCR_002380) (61) and Arabidopsis protein sequences from TAIR10 (RRID:SCR_004618) (97) were downloaded in fasta format on 4/17/21 and 2/26/21, respectively. Sequences were aligned using Exonerate v 2.2.0 (RRID:SCR_016088) (107) and the 5 best matches for each alignment were kept in the following format: --ryo `“>%qi length=%ql alnlen=%qal\n>%ti length=%tl alnlen=%tal\n”`. The resulting .gff2 was converted to a .gff3 using the script process_exonerate_gff3.pl (108), and the features ‘exonerate:protein2genome:local,’ ‘mRNA,’ and ‘CDS’ were changed to ‘protein2genome,’ ‘protein_match,’ and ‘match_part,’ respectively, for compatibility with MAKER.

### Running MAKER iteratively

MAKER was run similarly to (58,59), with the evidence detailed above and the custom repeat library created from the EDTA pipeline for masking (93). The output transcript and protein fasta files were extracted from MAKER’s first run and gene predictions with AED (Annotation Edit Distance) values <= 0.2 were used to train AUGUSTUS v 3.3.2 (RRID:SCR_008417) (109). Subsequently, MAKER was run a second time, with features from the first run’s .gff3 file being passed as hints to AUGUSTUS. After the run was complete, the resulting .gff3 and transcript and protein fasta files were again extracted as previously detailed (58).

### Polishing and filtering the annotation

Gene predictions output by MAKER’s second run were additionally processed to improve the annotation. First, the protein sequences were searched against the Pfam-A database (RRID:SCR_004726) (110) using hmmscan v 3.1b2 (RRID:SCR_005305) (111,112), and the predictions containing no known protein domains were removed. Second, defusion (60) was run to identify putatively-fused genes on the 24 chromosomes of the assembly (chr1[A, A’, B] – chr8[A, A’, B]). Defusion specializes in identifying potential tandem duplicates but does not typically identify fusions of genes with divergent intron-exon structures (chimeric fusions). However, it can extract and locally reannotate any sequences when given coordinates and breakpoint(s). Therefore, in addition to automatically identifying candidate gene fusions with defusion, we used an alternative method to identify candidates of the second class of gene fusions. Putatively fused loci from the initial MAKER gene set were identified when two or more distinct proteins-only gene predictions overlapped with gene predictions from the transcript plus protein MAKER run. (see Supplementary Materials, identify_fusion_candidates_w_PROTEIN_ONLY_datasets.bash). These candidate fusions were checked alongside those identified by defusion in IGV and break points were manually added to the .brk file as necessary (**Supplementary Figure 10**). The “defused” annotation was then filtered again to remove predictions lacking a Pfam domain (60). Lastly, putative DAM genes, *S*-RNases, and SFBs were manually annotated with Apollo v. 2.6.5 (RRID:SCR_001936) (64). The old gene models were then removed with agat (113) and replaced with the corrected models to produce the final annotation file.

### Assigning functions to genes

Functional information was assigned to the .gff and protein and transcript .fasta files via a BLAST+ v 2.9.0 (63) comparison of amino acid sequences to a Uniprot database (61) and several accessory scripts within MAKER (maker_functional_gff, maker_functional_fasta) (114). Moreover, Pfam (110), PANTHER (RRID:SCR_004869) (115), TIGRFAM (RRID:SCR_005493) (116), InterProScan (RRID:SCR_005829) (117), and Gene Ontology (GO) (RRID:SCR_002811) (118,119) database reference numbers or IDs were added to the .gff file by scanning the amino acid sequences with InterProScan and using the MAKER accessory script ipr_update_gff (114). Only hits with p-values < 1.0 × 10^-10^ were kept. Pfam, PANTHER, and TIGRFAM hits are also provided as separate .csv files that include the gene ID and functional descriptions.

### Assembly of a Prunus fruticosa draft genome

A similar approach to the *P. cerasus* ‘Montmorency’ genome was taken to assemble a draft genome of *P. fruticosa* but with some differences. Illumina gDNA reads’ 25-mers were counted using Jellyfish v 2.2.10 (79) and the resulting histogram was visualized using GenomeScope v2.0 (80). The estimated heterozygosity for *P. fruticosa*, which is also an obligate-outcrossing tetraploid, was 3.956%. The draft genome was created for the sole purposes of identifying *P. fruticosa*-like regions of the *P. cerasus* ‘Montmorency’ genome and estimating a divergence date between the two species via ortholog analyses. Because it is unknown whether *P. fruticosa* tetraploidy resulted from genome-doubling or interspecific hybridization (auto-vs allopolyploidization), parameters in Canu v 1.9 (81) were set similarly to those used for *P. cerasus* ‘Montmorency’ so that multiple haplotypes could be assembled from the PacBio reads. This provided assurance that if *P. fruticosa* comprises two different ancestral genomes, alleles from both subgenomes would most likely be assembled and included in ortholog analyses. The draft assembly was polished with the Illumina gDNA reads iteratively 3 times with Pilon v1.23 (82). A BUSCO analysis (43) was done to assess gene space, and Merqury (42) was used to determine phasing quality and genome-completeness.

### Annotation of a Prunus fruticosa draft genome

The polished *P. fruticosa* contigs were annotated using a similar pipeline to the one described for *P. cerasus* ‘Montmorency’. The RNAseq reads from *P. fruticosa* leaves, flowers, and developing fruits at 3 stages were processed in an identical manner to the *P. cerasus* RNAseq reads to produce a .gff3 file as input for MAKER. Uniprot and TAIR10 protein databases were aligned to the *P. fruticosa* contigs and processed similarly as well. Additionally, predicted proteins from *P. cerasus* ‘Montmorency’ with AED values < 0.3 were also used as evidence for MAKER. Gene finders SNAP (120) and AUGUSTUS (109) were trained on the .gff3 from the first MAKER run before using them for the second run. The final predicted gene set was filtered to keep predictions with known Pfam domains but lacking known TE domains. Due to limited resources, no manual annotation was performed on the *P. fruticosa* contigs. Assigning gene function to *P. fruticosa* gene predictions was done similarly as the *P. cerasus* ‘Montmorency’ annotation.

### Syntenic comparison of the Prunus cerasus ‘Montmorency’ assembly with P. persica

A synteny analysis was conducted between the 24 superscaffolds (chromosomes) of the *P. cerasus* ‘Montmorency’ assembly and a *P. persica* genome (46) using the SynMap tool within the Comparative Genomics Platform (CoGe) (45). Coding sequences (unmasked) of ‘Montmorency’ and *P. persica* were compared with default settings. Based on these synteny results (**Figure 1A**), k-mer clustering, and phylogenomic comparisons of syntelogs as described below, the 24 superscaffolds of the *P. cerasus* ‘Montmorency’ were named chr1[A, A’, B] – chr8[A, A’, B].

### Syntenic comparison of the Prunus cerasus ‘Montmorency’ assembly with representative progenitor genomes

Macrosyntenic comparisons of ‘Montmorency’ with the *P. avium* ‘Tieton’ v2.0 genome, the *P. fruticosa* draft genome, and itself were done using the MCScan package (RRID:SCR_017650) (121) from JCVI (RRID:SCR_021641) (121,122). We first built .cds and .bed files for each genome from the coding sequence and .gff files, respectively, then used command “jcvi.compara.catalog ortholog” to generate a list of syntenic blocks between either *P. avium* and ‘Montmorency’, *P. fruticosa* and ‘Montmorency’, or between ‘Montmorency’ and itself. All karyotype figures were constructed with the “jcvi.graphics.karyotype” command.

### k-mer clustering

We used Jellyfish 2.2.10 (79) to count 25-mers on each chromosome of ‘Montmorency’. 25-mers with fewer than 10 occurrences per chromosome were removed from the dataset and files were imported into R 4.2.0 (RRID:SCR_001905) (123). Further filtering was done if the 25-mers were not 1) present at 2x or more abundance in a homoeolog than in one of its sisters, and 2) present on all 24 chromosomes. We used the R function hclust() method “complete” to hierarchically cluster the 25-mers in the 24 and 22 chromosomes (excluding chromosome 8A and 8A’) and to construct dendrograms, and the package ‘pheatmap’ to create heat maps (RRID:SCR_016418). For the 25-mer clustering analysis to differentiate the A and A’ subgenomes only, the analysis was completed as described above with only 14 chromosomes (1[A, A’] - 7[A, A’]).

25-mer density per 1 Megabase (Mb) window of each chromosome was calculated as follows: (Number of group “X” 25-mers in a 1 Mb window × 25 bp) / (1 Mb). This is equivalent to the proportion of bases occupied by group “X” 25-mers in each 1 Mb window. Chromosome plots of 25-mer group densities were made in ggplot2 (RRID:SCR_014601) (124).

### Assessing read depth of the ‘Montmorency’ subgenomes

Read depth per position was assessed by mapping the ‘Montmorency’ Illumina reads to the assembly with Bowtie2 v. 2.3.4.2 default settings (83). SAMtools v 1.9 (RRID:SCR_002105) (100) was then used to sort and calculate depth at every position. From there, subgenomes were separated and concatenated end-to-end from chr1[A, A’, B] to chr8[A, A’, B]. For data reduction purposes, the average read depth per 1000 sites was plotted along the length of each subgenome and the median read coverage of the full genome was overlaid on the data (**Supplementary Figure 9**). Plots were created with R v. 4.2.1 (123) and ggplot2 (124).

### Phylogenomic comparisons of syntelogs to identify progenitor relationships

We used syntelogs (syntenic orthologs) between ‘Montmorency’ and each progenitor to assign regions of the assembly as either *P. fruticosa-like* or *P. avium-like*. Peptide and coding sequences of *Malus × domestica* ‘Gala,’ (125), *P. persica* ‘Lovell’ (46), *P. avium* ‘Tieton’ (22), *P. fruticosa* from the present study, *P. cerasus* ‘Montmorency’ subA, subA’, and subB were either downloaded from GDR or generated as described above. Orthogroups (groups of orthologous genes) were identified between these 7 “species” using OrthoFinder v. 2.5.4 (RRID:SCR_017118) (62). Multiple sequence alignments (MSAs) of orthogroups were done within OrthoFinder using MAFFT v. 7.480 (RRID:SCR_011811) (126) with no alignment trimming (-z). Only orthogroups including all 7 species were used for downstream analysis. Protein sequence alignments were converted to nucleotide alignments using PAL2NAL v. 14.1 (127), and raw cds alignments were trimmed with trimAl v. 1.4.1 (RRID:SCR_017334) (128)using flag `-automated1`. Alignments before and after trimming were visualized with MView (129) to ensure high quality of the resulting alignments. A phylogenetic tree for each orthogroup was created with RAxML-NG v 1.0.0 (RRID:SCR_022066) (130) using the gamma + GTR model, 500 bootstrap replications, and an apple ortholog outgroup. A two-column list of ‘Montmorency’ orthologs was used as input for PhyDS (131) to identify gene-gene sister relationships to either a *P. fruticosa* or *P. avium* ortholog with bootstrap values (BSV) of at least 80%. PhyDS requires a 2-column paralog list to extract relationships from phylogenetic trees, but no combination of two genes should be listed more than once. Thus, we extracted the IDs of all ‘Montmorency genes’ in all orthogroups and simply duplicated this list for the second column. We manually examined trees with a phylogenetic tree viewer (132) to ensure the paralog list and phyDS scripts were behaving as expected and extracted relationships where a ‘Montmorency’ ortholog was sister to a single representative progenitor gene (BSV >= 80%) using basic Unix commands (see Supplementary Materials). At the same time, syntenic gene pairs between A, A’, and B versus each representative progenitor (a total of 6 comparisons) were identified using the default settings of the python version of MCScan (121). The orthologous relationships identified with Orthofinder that fit the above criteria were interjoined with the syntenic orthologous relationships identified by MCScan using R version 4.2.1 (123). These high-confidence syntenic orthologs were mapped back to the ‘Montmorency’ assembly and labeled as either *‘P. avium-like’* or *‘P. fruticosa-like.’* The R package chromoMap (133) was used to visualize these results.

### Estimating divergence time of P. cerasus ‘Montmorency’ subgenomes from representative progenitor species

Separating A, A’, and B for ortholog analyses was done to ensure more single-copy orthologous groups could be identified between all “species” and used for phylogenomic dating. RAxML-NG was used to create phylogenetic trees for single-copy orthogroups as described above. Based on current knowledge of Rosaceae phylogenetics (1–3,66), the topolog(ies) for most accurately estimating the divergence of ‘Montmorency’ subgenomes and its representative progenitors from their most recent common ancestor (MRCA) should place apple as the super outgroup and peach as sister to the cherry lineage. Additionally, to calculate divergence time without error from possible homoeologous recombination, only orthogroups where ‘Montmorency’ homoeologs from each subgenome were predominantly sister to one progenitor over the other (*P. avium* or *P. fruticosa*) were included in the r8s analysis. This assumed that any previous homoeologous recombination taking place between the subgenomes did not replace >50% of the original sequence contributed by the progenitor. As a result, in any given tree, one homoeolog would not be able to pair with *P. avium* or *P. fruticosa* (i.e., there are 2 representative progenitors but 3 ‘Montmorency’ subgenomes). Thus, we were prepared to calculate node ages of multiple topologies to obtain MRCA divergence estimates for each subgenome and its most closely-related progenitor. This required each single-copy orthologous gene tree (n=336) to be manually inspected and the frequencies of each topology noted. The two most frequent topologies (**Supplementary Figure 17**) made it clear which progenitor subgenome A, A’, and B was most related to: subgenome B orthologs were almost always sister to *P. avium* while A and A’ orthologs were nearly equally likely to be sister to *P. fruticosa*. Orthogroup sequence alignments showing the two most frequent topologies were separately concatenated for all “species” (n=7) regardless of bootstrap support. Phylogenetic trees for the two concatenated alignments were created with RAxML-NG (130) as described above. Bootstrap replications (500 per topology) were used to calculate node age estimates and confidence intervals with r8s (RRID:SCR_021161) (134) and R v. 4.2.1 (123). Based on Xiang et al. 2017, the *Malus/Prunus* node was fixed at 95 million years ago (mya) and the peach/cherry node was constrained to a minimum age of 10 mya for both analyses. The smoothing parameter was set to 1, rate was set to gamma, and divergence time was set to penalized likelihood with the TN algorithm. 128633 sites were used to determine divergence times of the topology in **Supplementary Figure 17A** and 78,183 sites were used for the topology in **Supplementary Figure 17B**.

### Identification of the DAM genes and S-alleles

DAM (Dormancy-Associated MADS-box) genes were identified in ‘Montmorency’ and the *P. fruticosa* contigs with BLAST+ v. 2.9.0 (63), using *P. persica* DAM1 – DAM6 coding sequences from NCBI’s GenBank (RRID:SCR_002760) (135) as query and genomic sequence or transcripts from the MAKER pipeline as the target. The sequence IDs of the *P. persica* DAMs used as query were: DQ863253.2, DQ863254.1, DQ863256.1, DQ863250.1, AB932551.1, AB932552.1. Only matches with p-value < 1.0e-10 were kept with a max of 24 matches per query. This BLAST+ analysis identified DAM candidates in ‘Montmorency’ on chr1A, chr1A’, chr1B, and unanchored scaffold 2998, and in *P. fruticosa* on 7 different contigs, some containing only partial haplotypes of the expected 6 tandem genes with occasional erroneous fusions. Only genomic regions containing full haplotypes (6 tandem DAM genes) were manually annotated using Apollo. Chr1A, chr1A’, and chr1B in ‘Montmorency’ contained a full haplotype each, and two large contigs (8 and 33) contained a full haplotype each in *P. fruticosa*. Only these genes were used for phylogenetic comparisons with DAMs in other *Prunus* sp. These regions were confirmed to be syntenic between ‘Montmorency’ and *P. fruticosa*, and ‘Montmorency’ and *P. persica* (**Figure 5A, Figure 1A**).

*S*-alleles (*S*-RNase linked with an F-box protein/SFB) were identified in ‘Montmorency’ and *P. fruticosa* similarly to the DAMs with BLAST+ v. 2.9.0 (63) and the following complete cds sequences from NCBI’s GenBank (135) were used as query: P.cerasus*S*_36b_-RNase, P.cerasus*S*_36b3_-RNase, P.cerasus*S*_36b2_-RNase, P.avium*S*_6_-RNase, P.cerasus*S*_35_-RNase, P.cerasus*S*_36a_-RNase, P.cerasus*S*_13m_-RNase, P. cerasusSFB_36b_, P.cerasusSFB_36b3_, P.cerasusSFB_36b2_, P.aviumSFB_13_, P.cerasusSFB_35_, and P.cerasusSFB_36a_ (**Supplementary Table 5**). To be considered a full allele, an *S*-RNase and SFB with >90% identity to a query sequence had to be tightly linked (<100 kb apart).

### DAM phylogenetic comparisons

Protein sequences for *Arabidopsis thaliana* SEP3 and the six *P. avium* DAM genes were downloaded from NCBI **(Supplementary Table 4)**. Protein alignments were done with MUSCLE/3.8.31 (RRID:SCR_011812) (136) using default settings and the phylogenetic tree was constructed using RAxML-NG/1.0.0 (130). We used the PROTGTR+G model to infer the best maximum likelihood (ML) tree and mapped 500 bootstrap replicates onto the best ML tree to create the final phylogeny.

### S-RNase and SFB alignments

We downloaded the protein and coding sequences for the ‘Montmorency’ *S*-haplotypes from NCBI (RRID:SCR_006472) **(Supplementary Table 5)**. *P. fruticosa S*-alleles were aligned to their ‘Montmorency’ counterparts with MUSCLE/3.8.31 and alignment figures were made using the R package gggenes (**Supplementary Figure 16**) (124).

## Supporting information

All Supplementary Information

## Declarations

### Ethics approval and consent to participate

Not applicable.

### Consent for publication

Not applicable.

### Availability of data and materials

The datasets supporting the conclusions of this article are available in the Genomic Database for Rosaceae (GDR) https://www.rosaceae.org/ and NCBI’s Sequence Retrieval Archive (SRA) under BioProject number PRJNA922242. Scripts and example files associated with genome assembly, annotation, subgenome assignment analyses, and r8s estimates can be found at https://github.com/goeckeritz/Montmorency_genome. Scripts for k-mer hierarchal clustering, synteny, DAM gene phylogenetic analyses, and *S*-allele alignments can be found at https://github.com/KEBRhoades/Montmorency_genome.

### Authors’ contributions

CAH, AFI, and RV conceptualized the experiments. CZG performed genome assembly, annotation, subgenome assignment using orthologs, and divergence time estimate analyses. KER performed the k-mer hierarchal clustering, synteny, DAM gene phylogenetic analyses, and *S*-allele alignments. KLC provided expertise and assistance with annotation and contributed code. CZG, KER, and AFI wrote the manuscript. All authors assisted with editing the manuscript.

## Acknowledgements

We thank Dr. Shujun Ou for his assistance in running EDTA, and all members of the Aiden Lab and Dr. Ching Man Wai for their help in operating Juicer and 3D DNA. We are also grateful to Dr. Jose Ramon Planta and Dr. Jie Wang for their help with defusion, and to Dr. Nathan Dunn and Dr. Garrett Stevens for guidance with installing and navigating Apollo. Finally, we thank Dr. Pat Edger for his advice on all phylogenomic analyses.

