## Supplementary material for "A chromosome-scale assembly for tetraploid sour cherry (*Prunus cerasus* L.) ‘Montmorency’ identifies three distinct ancestral *Prunus* genomes": All Supplementary Information

*Supplementary Figures*

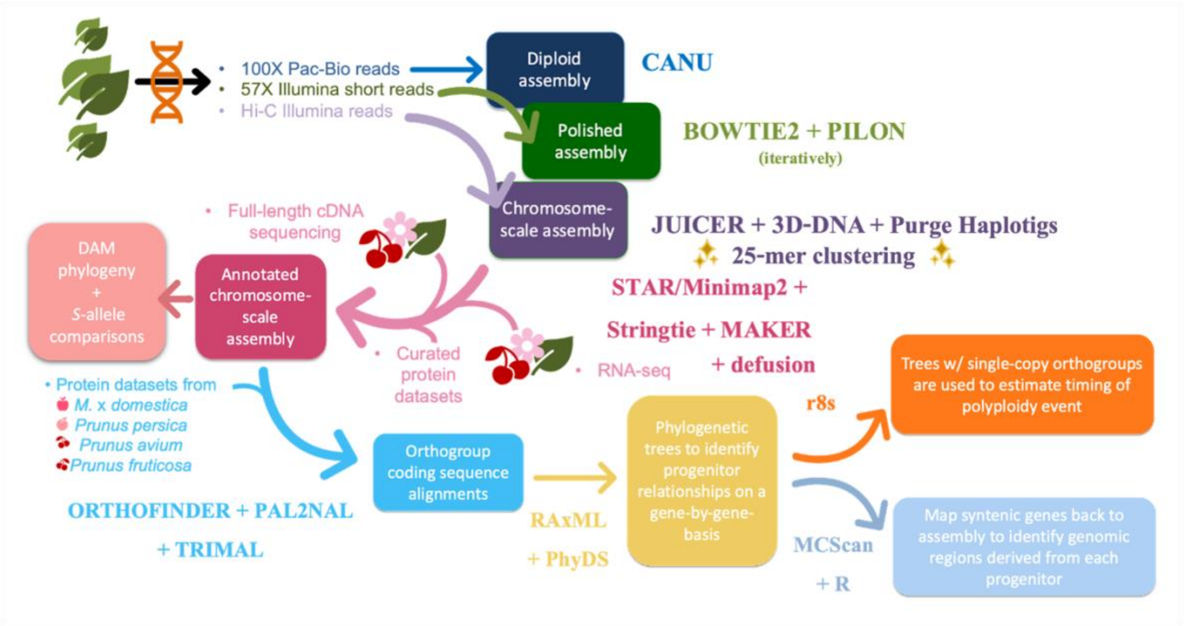


**Supplementary Figure 1**: **Workflow for the assembly and annotation of the *P. cerasus* 'Montmorency' genome and subsequent analyses.** Three types of DNA libraries were used to construct the 'Montmorency' reference genome. PacBio long-reads were assembled in Canu (81), and Illumina short-reads were used to polish this assembly with Pilon (82). Juicer + 3D-DNA (86, 87) were used to scaffold this polished assembly with Illumina short-reads from a Hi-C library. After initial scaffolding results and phylogenomic assessments, it was suspected the *Prunus avium*-like subgenome was likely in 2x dosage compared to the other two haplotypes and much more fragmented due to collapsing sequences during assembly. To reduce the complexity of the assembly and improve the representation of the *P. avium*-like subgenome, we used Purge Haplotigs (44) to set aside alternative alleles, hoping that soft purging (>99% alignment) would predominantly remove alleles from this subgenome and not from the other two. From the initial scaffolding results, we knew this subgenome was far more fragmented; therefore, contigs of size greater or equal to 400 kb were added back into the assembly after purging and prior to scaffolding. Once the 24 linkage groups were scaffolded and affirmed to be near-complete genomes [based on low BUSCO (43) duplication and assessment of k-mer spectra in Merqury(42)], we used k-mer clustering to identify 25-mers distinguishing 3 *Prunus* genomes, with the exclusion of two homoeologs from chromosome set 8. High-quality transcripts from various tissues alongside curated protein datasets provided evidence for gene prediction with MAKER (59). Additional curation of the annotation was done using defusion and Apollo (64). With the annotated 'Montmorency' sequence, the DAM and *S*-allele genes were compared to representative progenitors. Additionally, orthogroup relationships allowed for the estimation of each subgenome's divergence date from its representative progenitor with single-copy orthogroups (134). Finally, all orthologous groups were examined and relationships of each subgenome to a progenitor were extracted (131), checked for synteny (121), and mapped back to the scaffolded genome assembly to determine the genomic regions most probably derived from each representative progenitor.

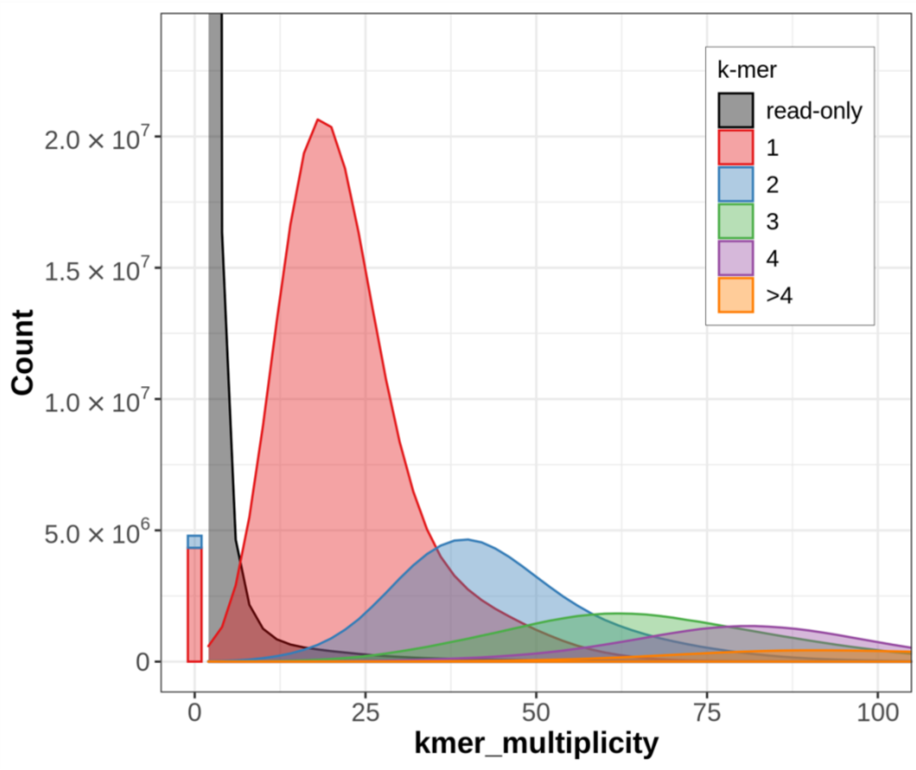


**Supplementary Figure 2: Merqury plot for the 'Montmorency' genome assembly.** The k-mer spectra of the Illumina reads illustrates the number of times a k-mer appears in the *P. cerasus* 'Montmorency' assembly. The red peak represents k-mers of size 25 bp found in the short-read dataset approximately 15-20X and once in the assembly, the blue peak represents 25-mers found in the Illumina dataset 30-40X and twice in the assembly, and so on. The concordance of the k-mers' multiplicity in the Illumina dataset with the number of times the kmer is found in the assembly indicates good haplotype phasing and the assembly of all 4 possible alleles in the *P. cerasus* 'Montmorency' assembly. The black portion of the spectra represent k-mers from short-reads not found in the assembly, and the small red and blue bar represents k-mers that are in the assembly but not in the short reads. These k-mers likely represent PacBio sequencing errors that failed to be corrected during polishing with Pilon.

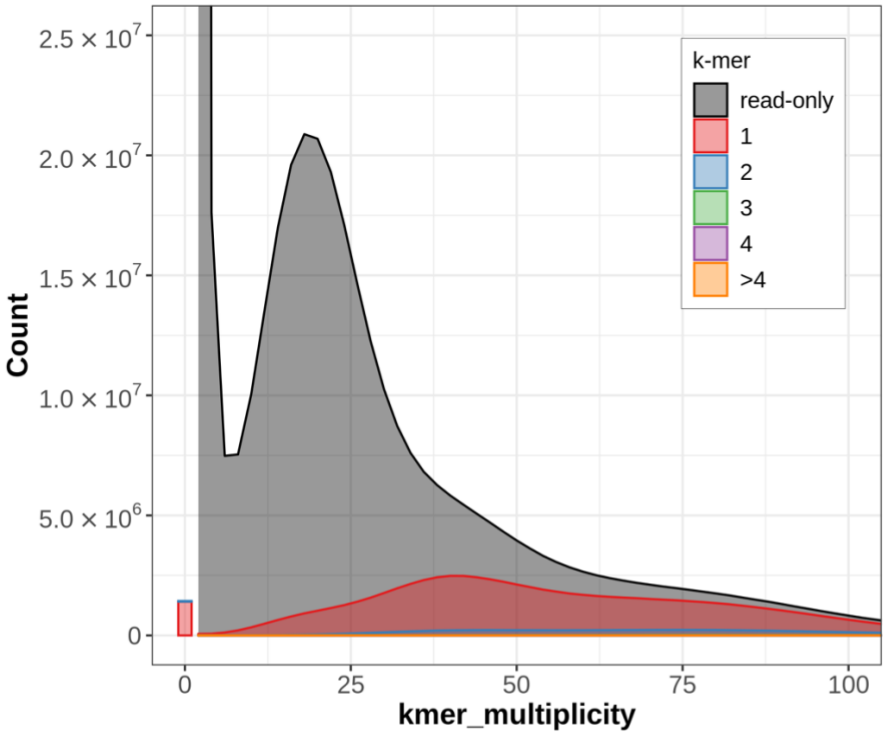


**Supplementary Figure 3: The purged portion of the 'Montmorency' assembly represents one of four alleles**. The black area represents k-mers found in the Illumina reads but not in this subset of the assembly. The red area indicates sequences of all multiplicities (i.e., in the 'Montmorency' genome one, two, three, or four times) are present in this purged subset only once, meaning alleles from only one haplotype had been purged.

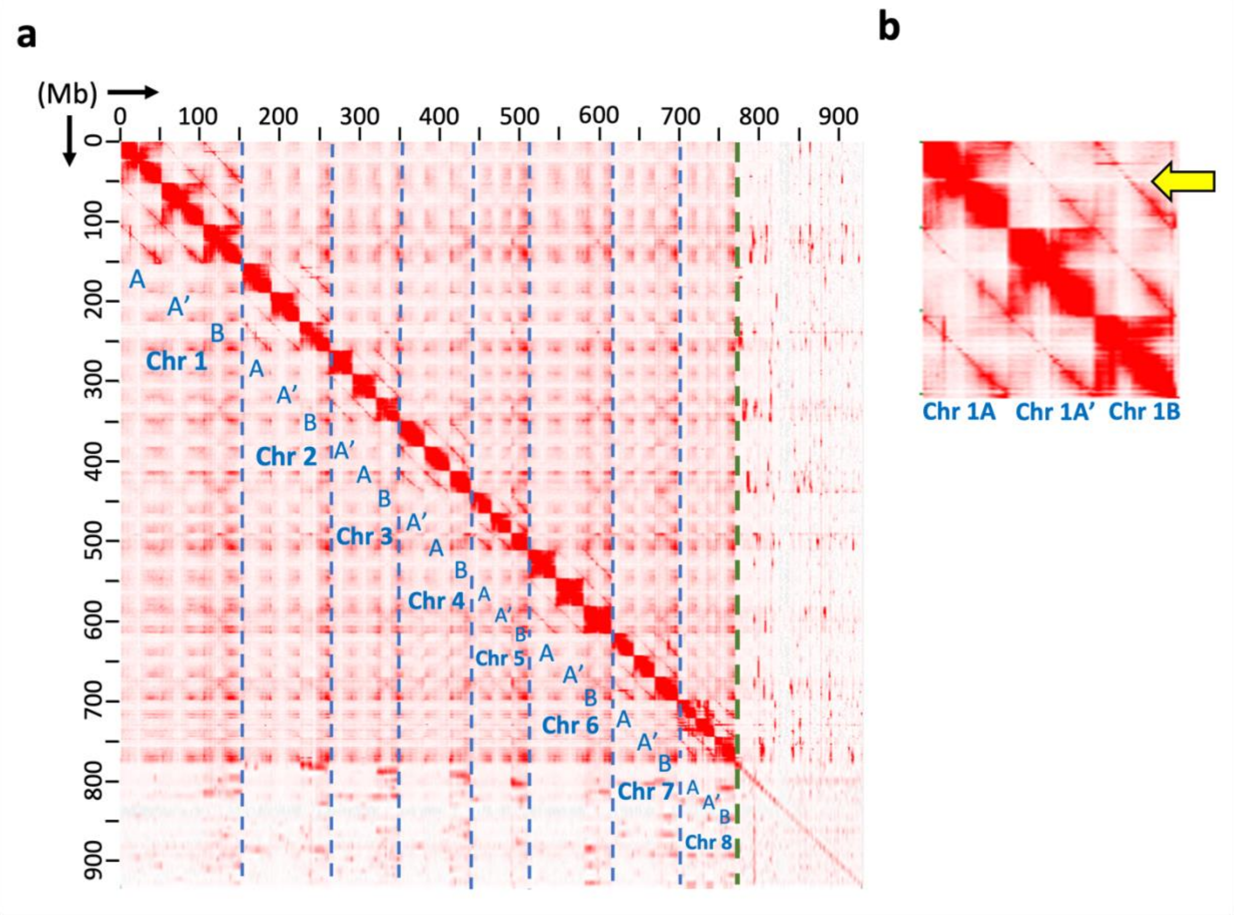


**Supplementary Figure 4**: **Hi-C matrix of the 'Montmorency' assembly indicates 24 linkage groups, with three homoeologs per set**. a) The dashed blue lines delineate the 8 ancestral groups of *Prunus*; 3 homeologs were assembled per group. The green dashed line separates the scaffolded portion of the assembly from unanchored contigs. Note that not all unanchored contigs are shown as some were removed prior to scaffolding to obtain a better representation of subgenome B. (see Methods) b) *Prunus* chromosome set 1 is shown for the 'Montmorency' assembly. A yellow arrow points to Hi-C interactions suggestive of a homoeologous relationship between chr 1A and chr 1B.

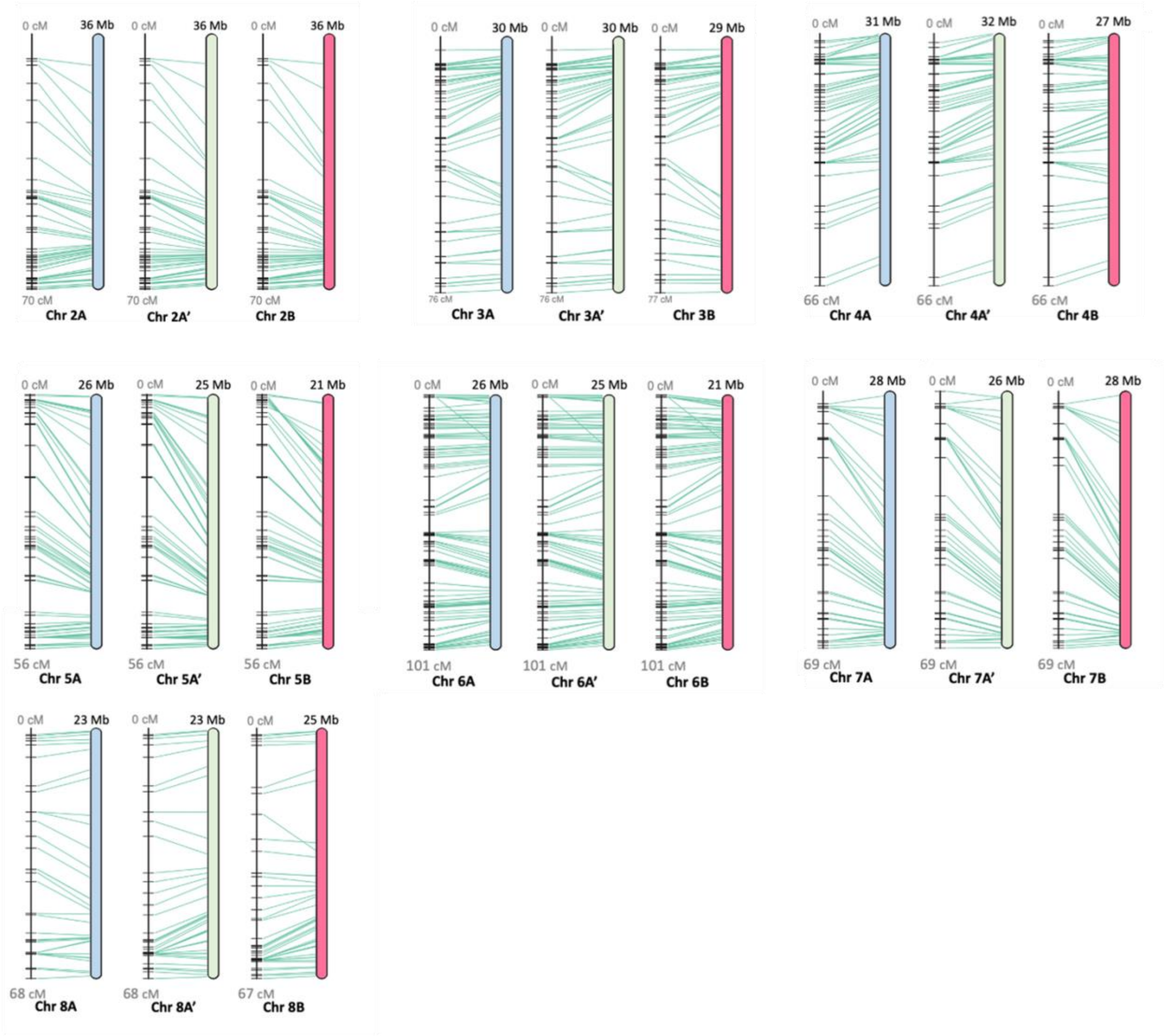


**Supplementary Figure 5: Linearity comparison of linkage groups 2-8 and a published sour cherry genetic map** (11). 545 Markers from an F1 sour cherry cross in which ‘Montmorency’ was the female parent were mapped to the assembly and the results display high collinearity between the linkage map and assembly. Green lines connect the markers on the genetic map (left) to the physical location in the assembly (right). Each horizontal black line on the genetic map represents one marker. Post-filtering, 426 of the 545 markers mapped exactly once to each subgenome. Chromosome sets from subgenome B are representative of two possible haplotypes. Linkage group 1 can be viewed in Figure 2. Figures were generated using ALLMAPS (96).

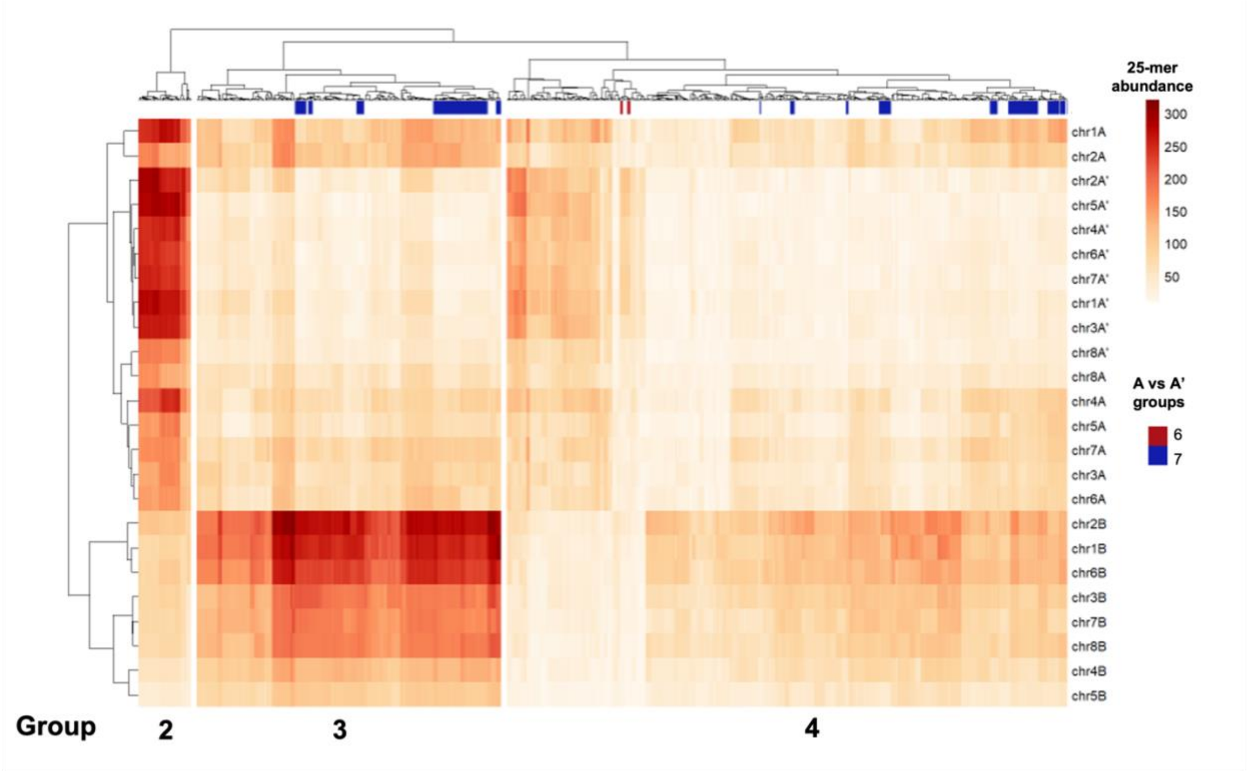


**Supplementary Figure 6**: **Heat map of the hierarchical clustering of 25-mer densities on all 24 'Montmorency' chromosomes**. The differential abundance of the 25-mers in Group 2 and Group 3 enable the clustering of 16 A/A' subgenome chromosomes from 8 B subgenome chromosomes. Group 1 is not shown in the heatmap because it only contains two 25-mers and heavily skews the normalization of the heat map. The 25-mers common to this analysis and the 25-mer analysis differentiating A and A' (excluding chromosomes 8A and 8A') are annotated according to colors used in Supplementary Figure 7. Group 5 25-mers are entirely unique to subgenomes A and A' and thus are not annotated here. Heat map color indicates the abundance of each 25-mer by chromosome.

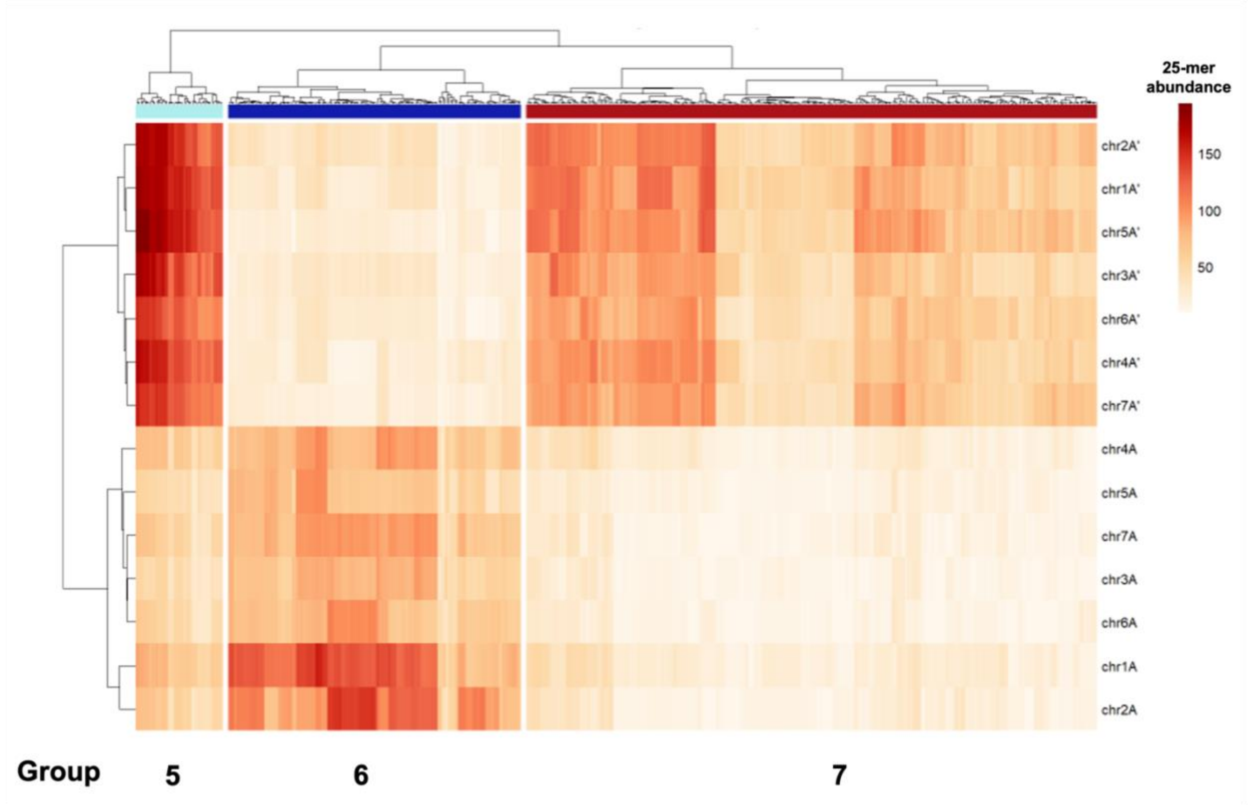


**Supplementary Figure 7**: **Heat map of the 25-mer densities corresponding to the hierarchical clustering of 14 chromosomes**. These 25-mer groups differentiate the seven chromosomes of subgenomes A and A’. The 14 chromosomes in this analysis are a subset of the 16 chromosomes that formed one clade in the original 24 chromosome analysis (Supplementary Figure 6), but here 8A and 8A’ are excluded. Group 5 is unique to subgenomes A and A’. The colored bars beneath the dendrogram on the x-axis are present for the purpose of indicating 25-mers common to this clustering analysis and the full 24 chromosome analysis shown in Supplementary Figure 6. Heat map color indicates the abundance of each 25-mer by chromosome.

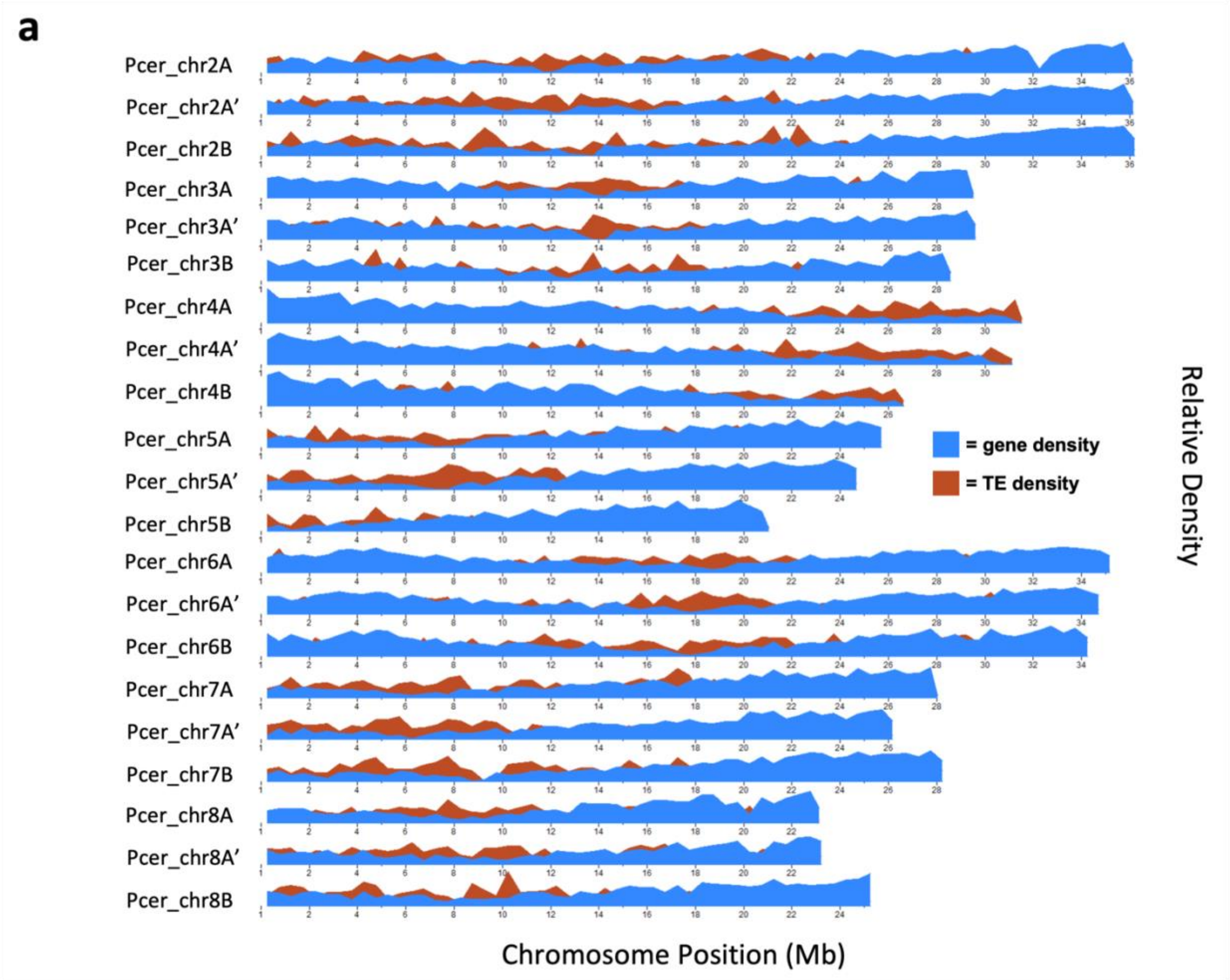
a
Pcer_chr2A
Pcer_chr2A'
Pcer_chr2B
Pcer_chr3A
Pcer_chr3A'
Pcer_chr3B
Pcer_chr4A
Pcer_chr4A'
Pcer_chr4B
Pcer_chr5A
Pcer_chr5A'
Pcer_chr5B
Pcer_chr6A
Pcer_chr6A'
Pcer_chr6B
Pcer_chr7A
Pcer_chr7A'
Pcer_chr7B
Pcer_chr8A
Pcer_chr8A'
Pcer_chr8B
= gene density
= TE density
Relative Density
Chromosome Position (Mb)

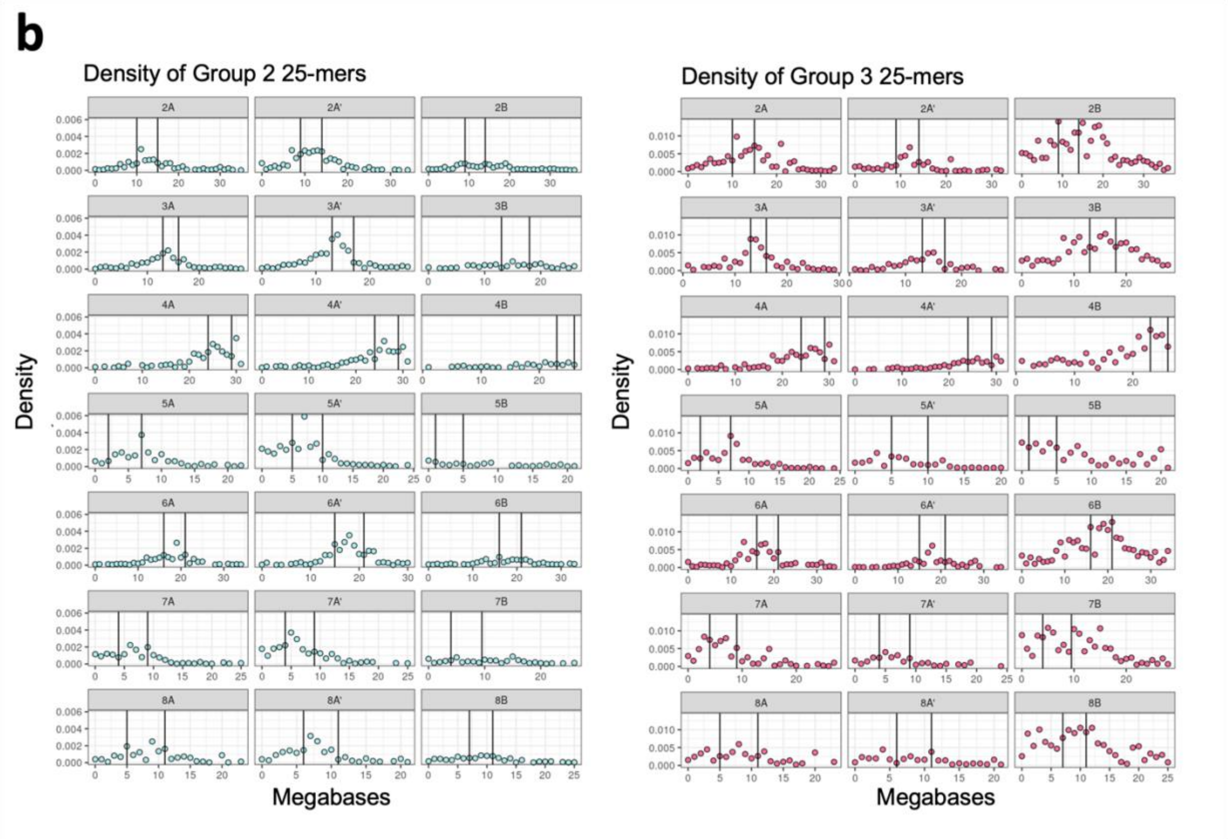
b
Density of Group 2 25-mers
Density of Group 3 25-mers
2A
2A'
2B
3A
3A'
3B
4A
4A'
4B
5A
5A'
5B
6A
6A'
6B
7A
7A'
7B
8A
8A'
8B
Density
Megabases

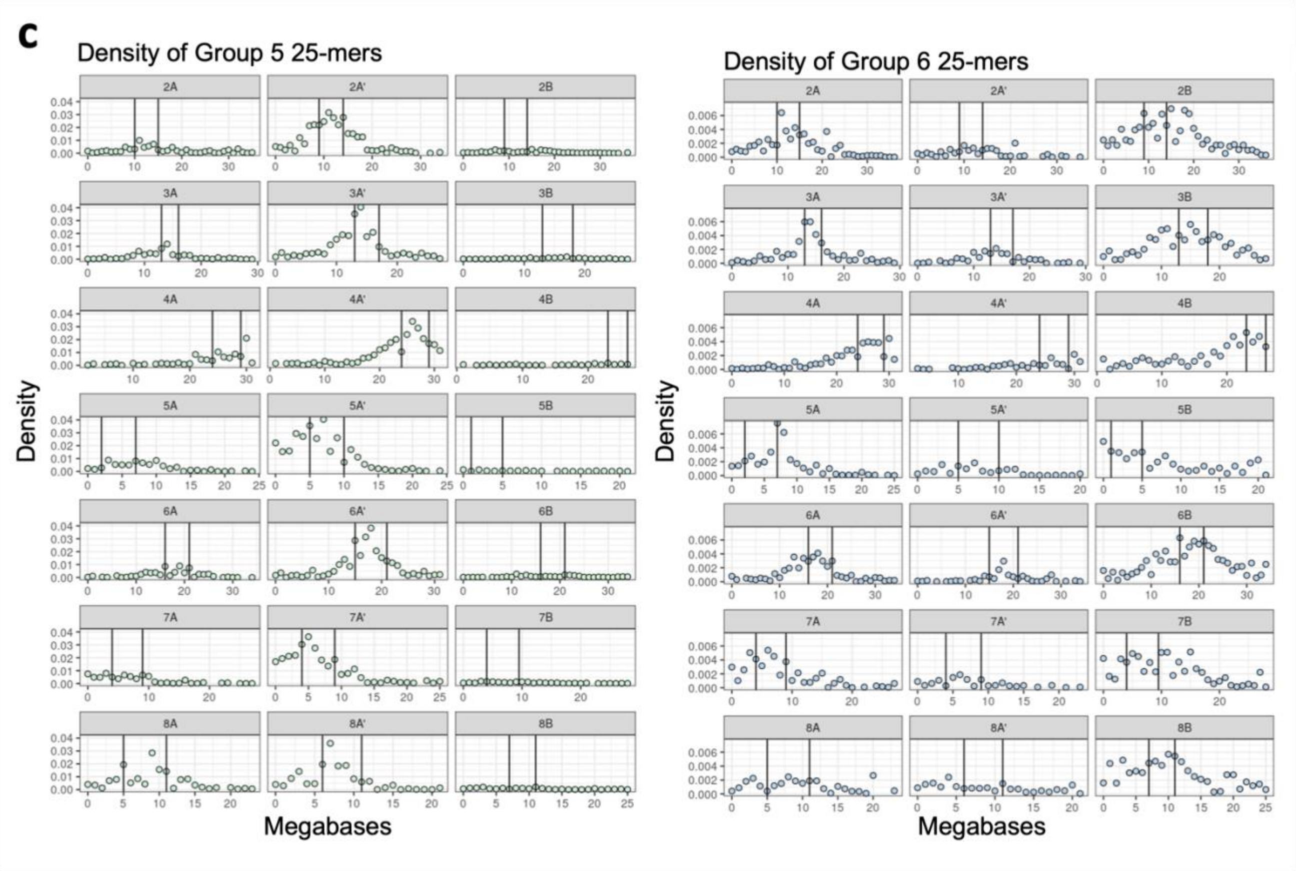


**Supplementary Figure 8: 25-mer group densities differentiating the 'Montmorency' subgenomes peak at approximate centromeres.** a) Gene and transposable element (TE) densities plotted along the lengths of chromosome sets 2 – 8. The centromeres were estimated to be the regions that coincide with relatively low gene and high TE densities. b) Group 2 and Group 3 25-mer densities (Supplementary Fig. 6) are plotted along the lengths of chromosome sets 2 – 8. These groups separate the A/A' subgenomes from B in the 24 chromosomes clustering analysis (Fig. 3a). c) Group 5 and Group 6 25-mer densities (Supplementary Fig. 7) plotted along the lengths of chromosome set 2 – 8. These distinguish A and A' from one another when only 14 chromosomes are included in the clustering. All 25-mers in Group 5 are unique to A and A' (note the flat 0.00 density for subgenome B). In both b) and c), vertical black lines mark approximate centromere locations based on low gene density and high TE density. See Figure 4 for corresponding data for chromosome set 1.

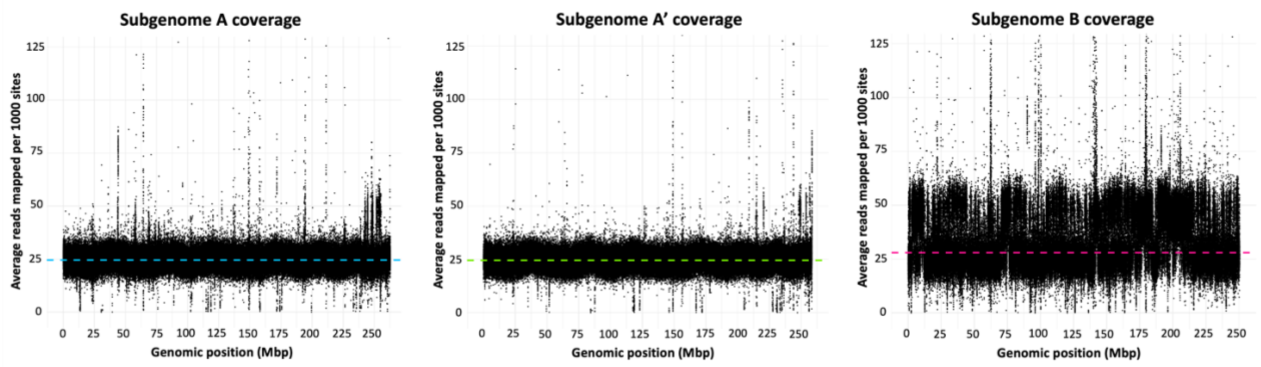


**Supplementary Figure 9: Illumina reads mapped back to the 'Montmorency' assembly show regions of subgenome B have approximately twice the coverage as A and A'**. Chromosomes were concatenated end-to-end for each subgenome, so the x-axis represents the entire length of each subgenome. The y-axis is average read depth per 1000 sites. The dashed line in each plot indicates median coverage of that subgenome. Spikes in coverage along the length of subgenome B suggest a higher copy number compared to subgenomes A and A'.

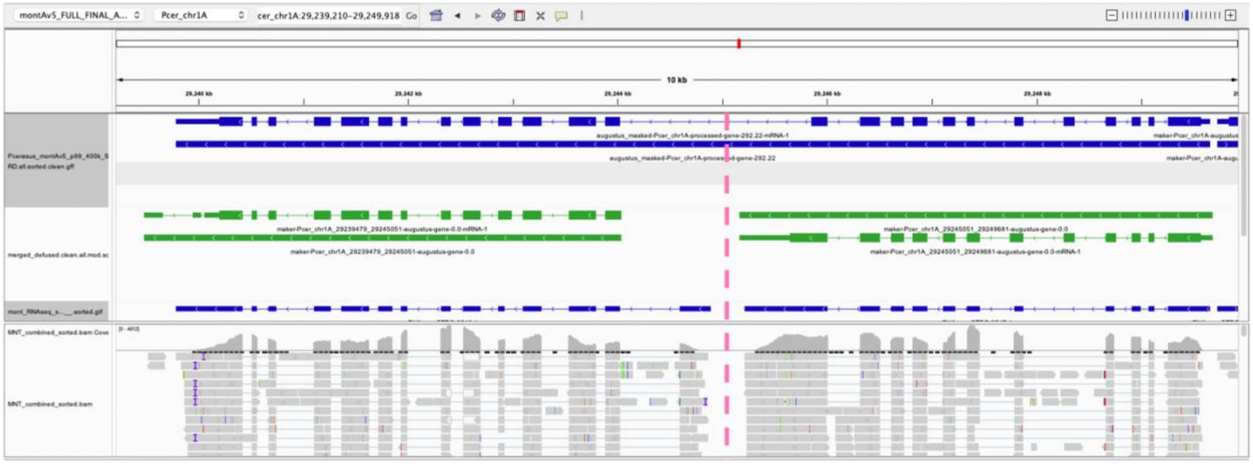


**Supplementary Figure 10: IGV screenshot illustrating how the defusion pipeline can improve gene model predictions**. The top track (blue) represents one large gene model right after running MAKER. However, according to the Stringtie2 (101) transcript assembly (track 3, blue) and RNAseq coverage and alignments (tracks 4 and 5 respectively, gray), this large gene model should be two, with an approximate gene boundary indicated by the dashed vertical pink line. Once defusion is provided with this boundary (breakpoint), it reruns MAKER locally within the two genomic regions. The second track (green) shows the original gene model properly became two and most exon / intron structures are accurate according to the evidence.

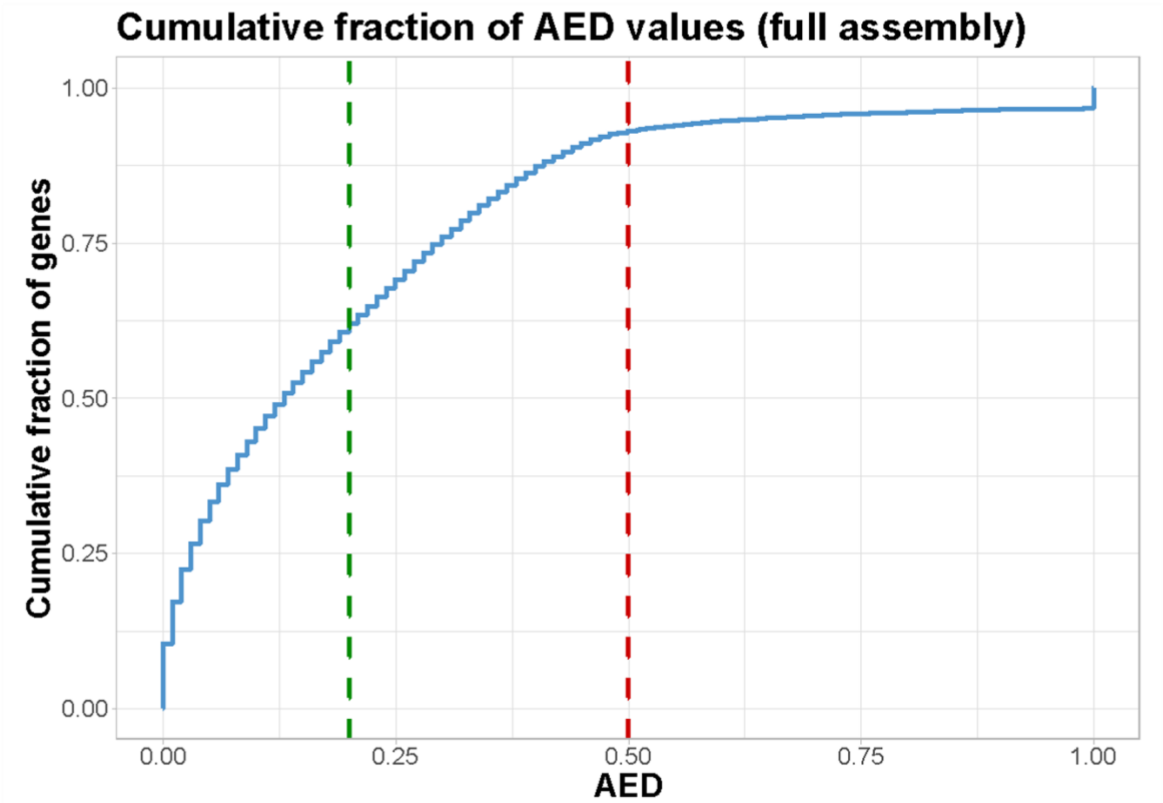


**Supplementary Figure 11**: **Cumulative fraction of Annotation Edit Distance (AED) values for gene predictions in the 'Montmorency' assembly**. The lower the AED (59) value for a gene, the more concordant it is with provided evidence (protein, RNA, etc). 0 indicates perfect concordance between a gene prediction and the evidence at that location, whereas 1 indicates no evidence supports a gene prediction. Dashed lines indicate the fraction of genes showing AED values less than or equal to 0.2, while the red dashed line indicates the fraction of genes showing AED values less than or equal to 0.5.

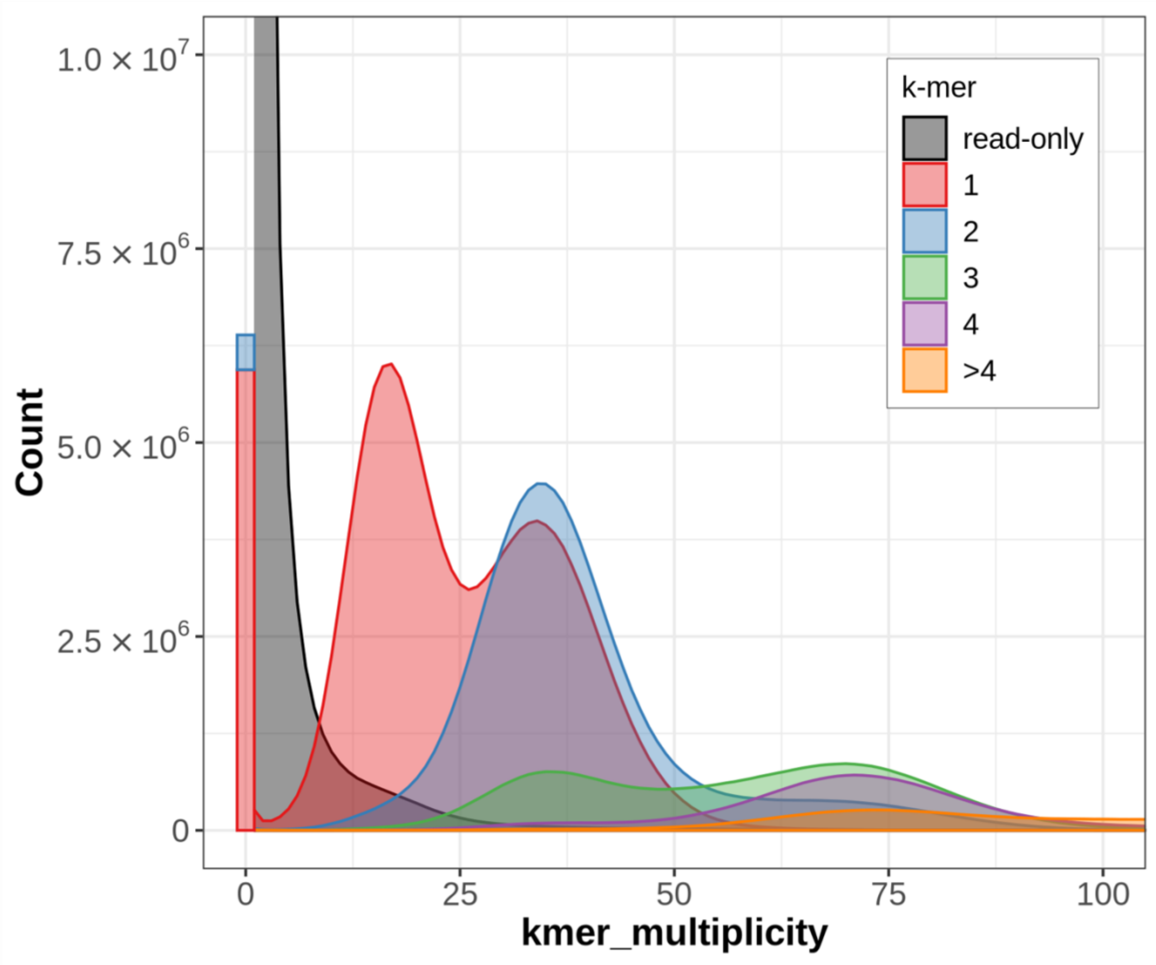


**Supplementary Figure 12**: **The k-mer spectra of the Illumina short-reads against the *Prunus fruticosa* draft assembly**. Color indicates the number of times a k-mer from the Illumina short-reads appears in the draft *P. fruticosa* assembly. The red peak represents kmers of size 25 bp found in the short-read dataset approximately 15 - 20X and once in the assembly, the blue peak represents 25-mers found in the Illumina dataset 30-40X and twice in the assembly, and so on. A second red peak for kmers found 30 - 40X in the short-read dataset indicates the presence of collapsed haplotypes, as their multiplicity (30 - 40X) indicates they should be found approximately twice in the assembly. The green peaks indicate regions of the genome that should be found in the assembly three times have been collapsed or artificially duplicated. The black portion of the spectra represent k-mers from short-reads not found in the assembly, and the red and blue bar represents k-mers that are in the assembly but not in the short reads. These k-mers likely represent PacBio sequencing errors that failed to be corrected during polishing with Pilon.

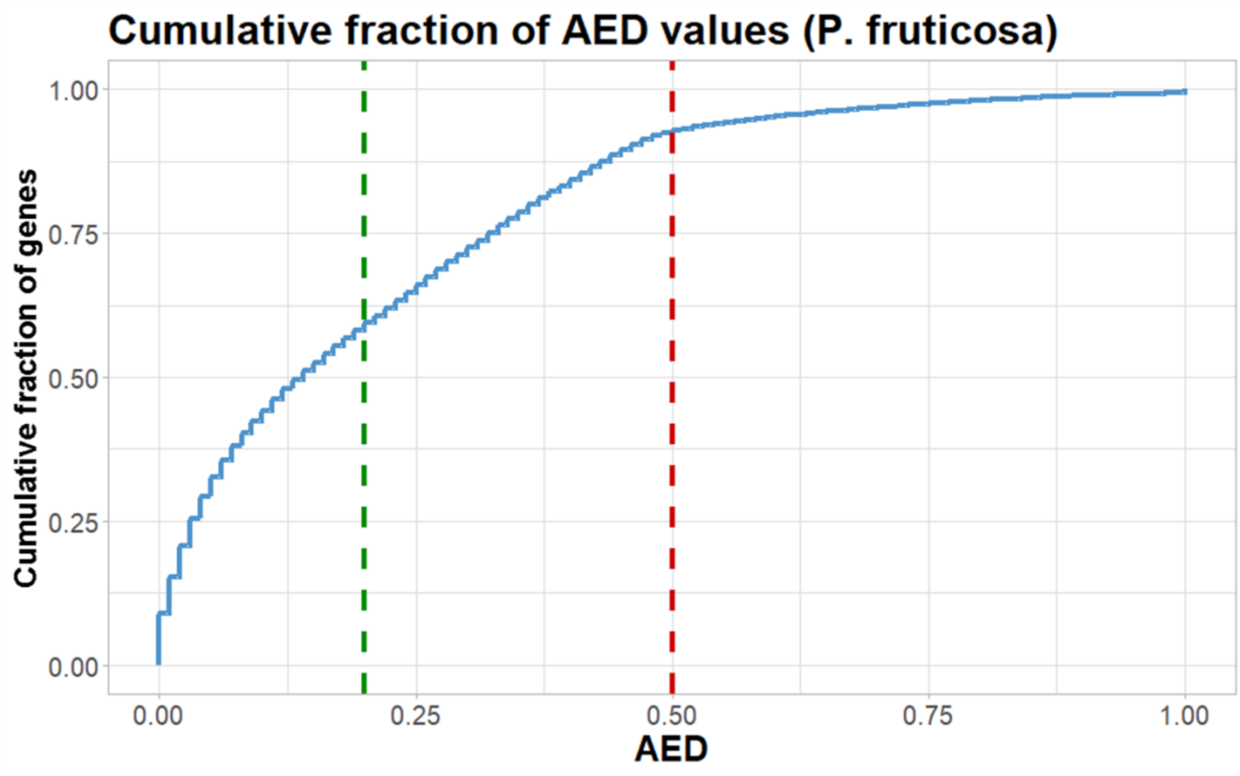


**Supplementary Figure 13**: **Cumulative fraction of Annotation Edit Distance (AED) values for gene predictions in the *Prunus fruticosa* draft assembly.** The lower the AED value for a gene, the more concordant it is with provided evidence (protein, RNA, etc). 0 indicates perfect concordance between a gene prediction and the evidence at that location, whereas 1 indicates no evidence supports a gene prediction. Dashed lines indicate the fraction of genes showing AED values less than or equal to 0.2, while the red dashed line indicates the fraction of genes showing AED values less than or equal to 0.5.

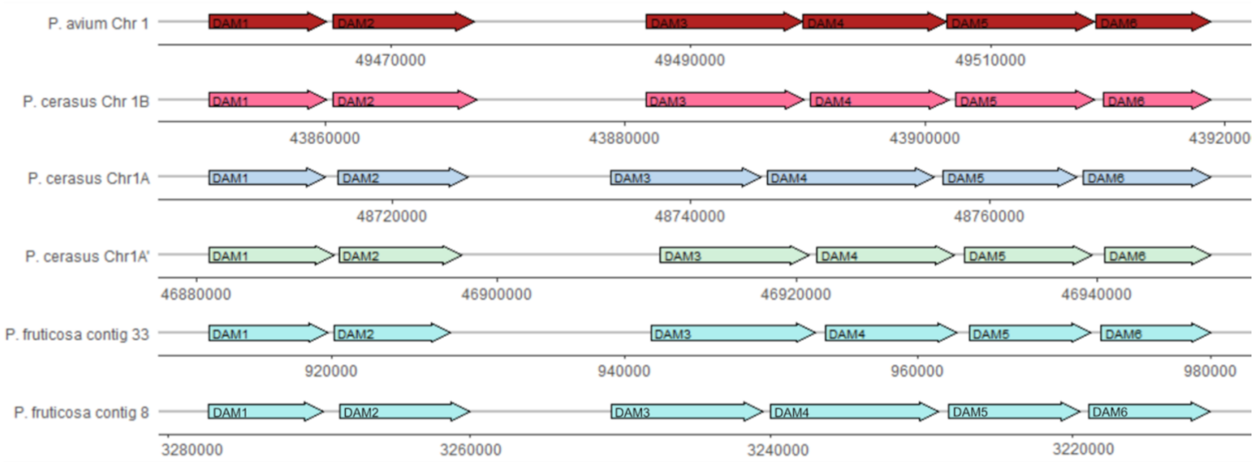


**Supplementary Figure 14: The DAM genes identified in 'Montmorency' and in *Prunus fruticosa* all show the *Prunus*-characteristic tandem arrangement**. In both 'Montmorency' and *P. avium* the Dormancy Associated MADS-box genes (DAMs) are at the bottom of chromosome 1. Contigs 8 and 33 in the *P. fruticosa* assembly are syntenic with these regions in 'Montmorency' (see Figure 6a). Note that contig 8 is in the reverse orientation relative to other contigs and chromosomes.

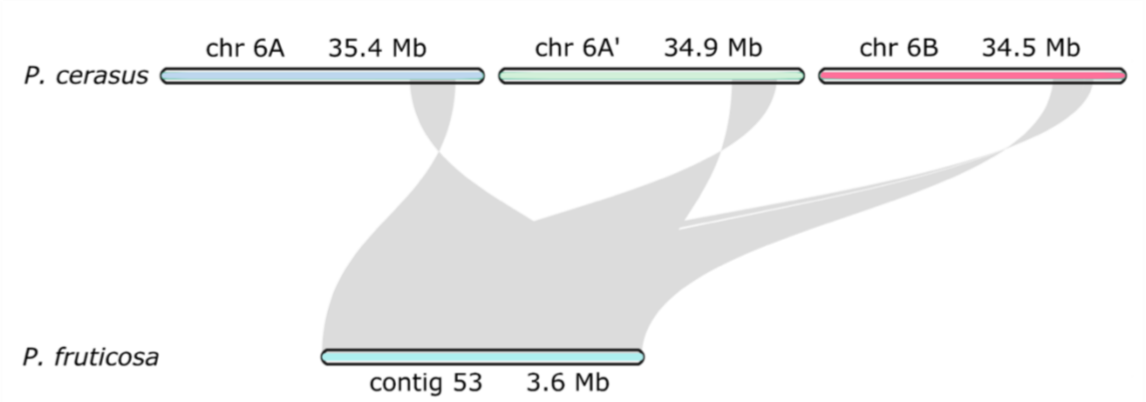


**Supplementary Figure 15**: **The region on 'Montmorency' chromosome 6 where the *S*-alleles were identified are highly syntenic with contig 53 in *P. fruticosa,* where one *S*-allele was identified**. Notice the contig is given in the reverse orientation relative to the 'Montmorency' chromosomes; this is not evidence of a true inversion.

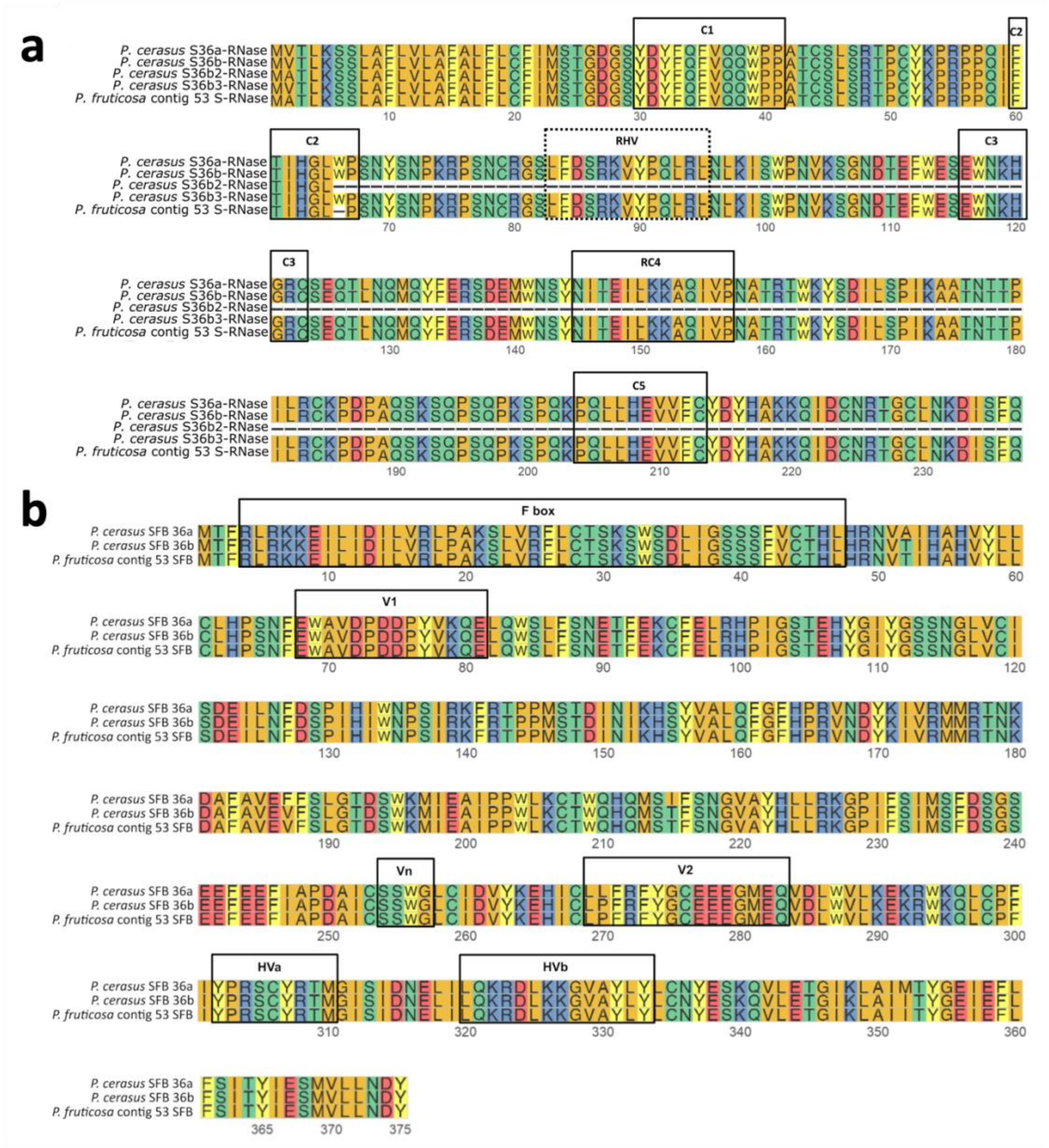


**Supplementary Figure 16**: **Amino acid alignments of *S*-allele components (*S*-RNase + *S*-locus F-box protein/SFB) between published 'Montmorency' *S*36 variants and the allele identified in *Prunus fruticosa* on contig 53.** a) *S*-RNase amino acid alignments. The five conserved regions [C1, C2, C3, RC4, and C5 (35)] are identified with solid boxes, and the hypervariable region [RHV (35)] is delineated with a dotted box; this region is unique to rosaceous *S*-RNases. The – in region C2 indicates where a premature stop codon is predicted to truncate the *S*-RNase in *P. fruticosa,* making the sequence 100% identical to *S*36b3; however, the rest of the protein is presented as if the stop codon were read through to indicate the presence of the other conserved regions. b) SFB amino acid alignments. Only SFB36a and SFB36b are shown as SFB36b2 and SFB36b3 are identical to SFB36b (40) Boxes identified the locations of the F-box motifs [V1, V2, Hva and HVB (33) and Vn (36)]. The SFB identified on *P. fruticosa* contig 53 is 100% identical to the published *P. cerasus* SFB 36b sequence.

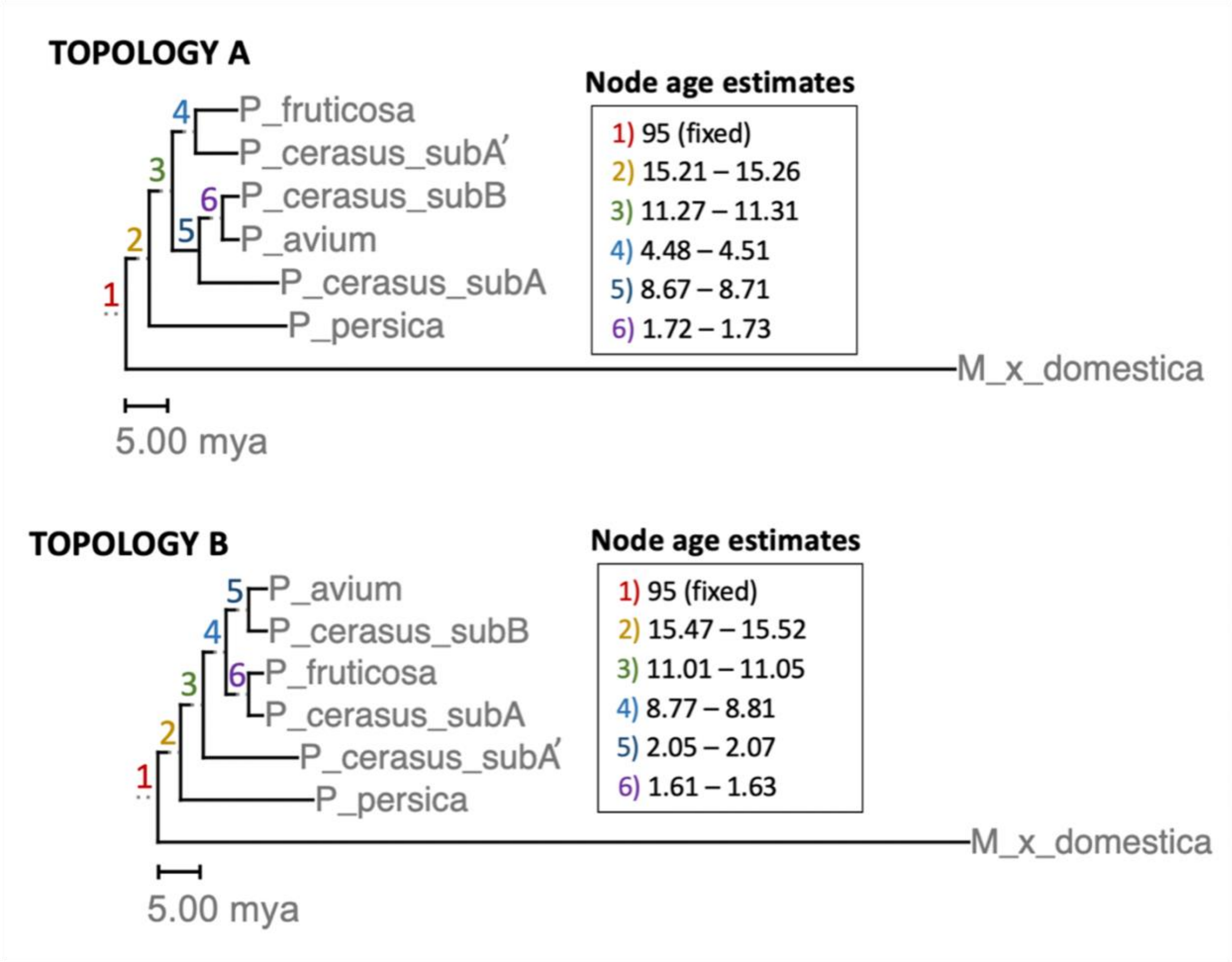


**Supplementary Figure 17**: **Chronograms of the two most frequent topologies produced from single-copy orthologous genes**. Nodes are numbered on the phylogeny. The 95% CI of node age estimates are given for each node on the right in millions of years ago (mya). The confidence intervals for each phylogeny's node ages were calculated based on 500 bootstrap replicates for each topology. In both analyses, Node 1 was fixed at 95 mya and Node 2 was constrained to be a minimum of 10 mya based on Xiang et al. 2017 (2). Chronograms were produced with r8s (134).

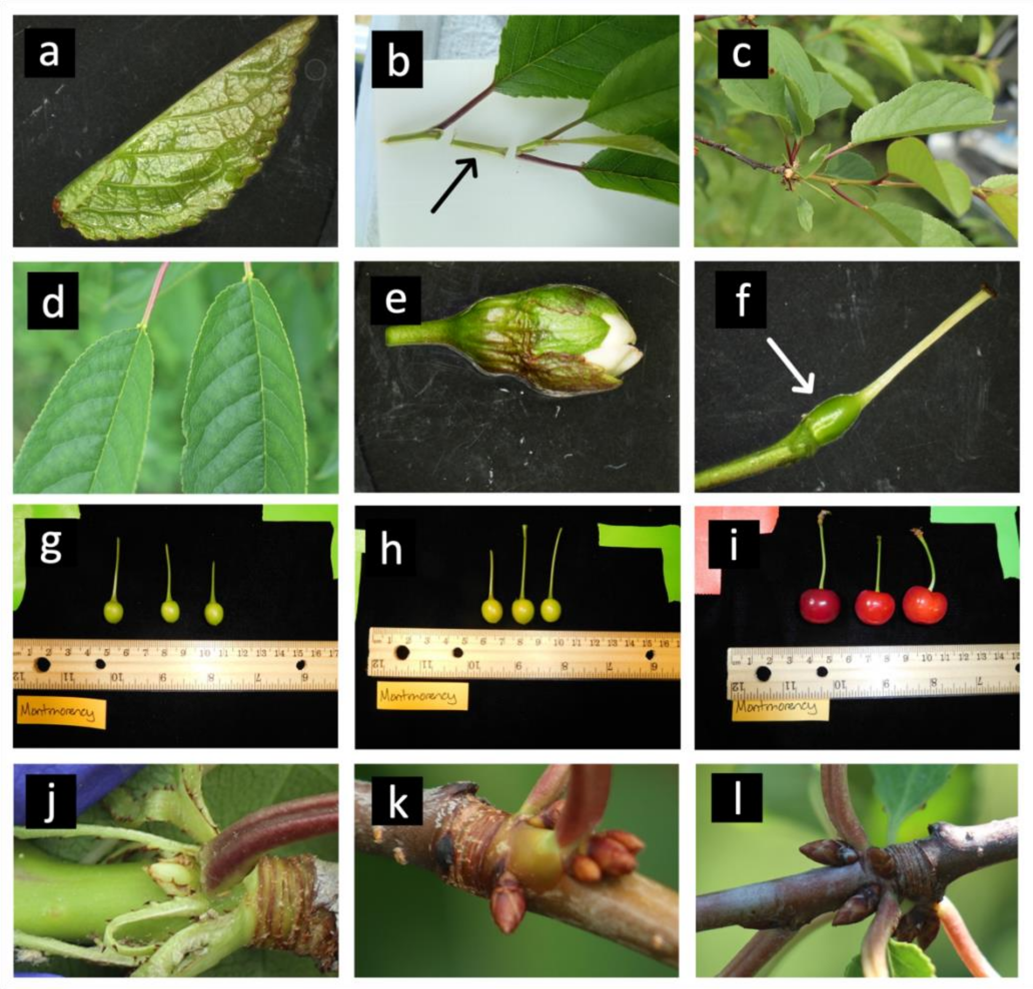


**Supplementary Figure 18**: **Photos of tissue collected for RNA extraction and annotation of the ‘Montmorency’ genome**. a) young leaf, b) internode of current year’s growth (black arrow), c) fully-expanded, mature leaves in late spring (6/12/19), d) fully expanded, mature leaves in midsummer (7/18/19), e) whole flower at balloon stage, f) fruit stage I (ovary post-pollination, white arrow), g) beginning of fruit stage II (5/28/19), h) end of fruit stage II (6/17/19), i) fruit stage III, j) vegetative meristem prior to floral initiation in late spring (6/12/19), k) meristem collected mid-summer at approximate time of floral initiation (7/18/19), l) floral buds during organ differentiation (8/22/19). Not individually pictured but also sequenced as separate libraries: stage III fruit exocarp (skin), stage III fruit mesocarp.

### Supplementary Table 1: Summary of the annotation metrics for the 'Montmorency' assembly

| | | | | | |
|---|---|---|---|---|---|
| **Protein-coding gene count** | subA | subA' | subB | Total (incl. unanchored) | |
| | 25272 | 24863 | 24914 | | 97254 |
| **AEDs summary** | min | median | mean | max | |
| | 0 | 0.13 | 0.1993 | 1 | |
| **Average protein-coding gene length** | **base pairs** | | | | 3296 |
| | **amino acids** | | | | 425.5 |
| **transcript BUSCO (viridiplantae db10)** | complete | singletons | duplicates | missing | fragmented |
| scaffolded assembly | 97.4% | 7.3% | 90.1% | 1.4% | 1.2% |
| subgenome A | 89.2% | 87.3% | 1.9% | 7.7% | 3.1% |
| subgenome A' | 90.4% | 88.0% | 2.4% | 4.9% | 4.7% |
| subgenome B | 87.8% | 84.5% | 3.3% | 8.4% | 3.8% |
| **protein BUSCO (viridiplantae db10)** | complete | singletons | duplicates | missing | fragmented |
| scaffolded assembly | 97.2% | 6.6% | 90.6% | 0.9% | 1.9% |
| subgenome A | 87.5% | 86.1% | 1.4% | 9.0% | 3.5% |
| subgenome A' | 90.1% | 88.2% | 1.9% | 4.7% | 5.2% |
| subgenome B | 86.3% | 84.2% | 2.1% | 8.8% | 4.9% |

AED = Annotation Edit Distance (reference 49); BUSCO = Benchmarking Universal Single-Copy Orthologs (reference 43)

**Supplementary Table 2: Summary of the *Prunus fruticosa* draft assembly metrics**

| | | | | | |
|---|---|---|---|---|---|
| Estimated haploid genome size (kmer analysis; k = 25) | | | | | 532 Mb |
| Estimated Heterozygosity (total) | | | | | 3.956% |
| class | aaab | aabb | aabc | abcd | |
| | 0.276% | 3.360% | 0.001% | 0.319% | |
| Assembly size | | | | | 986 Mb |
| NG50 | | | | | 5.76 Mb |
| Number of contigs | | | | | 3932 |
| Contig length stats (bp) | min | median | mean | max | |
| | 5389 | 75874 | 250776 | 18693262 | |
| BUSCO (viridiplantae db10) | complete | singletons | duplicates | missing | |
| | 99.10% | 6.40% | 92.70% | 0.70% | |
| Estimated % repeats | LTR | TIR | Helitron | Total | |
| | 35.45% | 11.15% | 1.33% | | 47.93% |
| LAI | | | | | 15.58 |

Mb = megabases; LG = Linkage Group; NG50 = 50% of the estimated genome size is contained in contigs of equal or greater value; BUSCO = Benchmarking Universal Single-Copy Orthologs (reference 43); LTR = Long Terminal Repeat; TIR = Terminal Inverted Repeat; LAI = LTR Assembly Index (reference 90).

**Supplementary Table 3: Summary of the annotation metrics of the *Prunus fruticosa* draft assembly**

| | | | | | |
|---|---|---|---|---|---|
| **Protein-coding genes** | | | | | 102379 |
| **AEDs summary** | min | median | mean | max | |
| | 0 | 0.14 | 0.20 | 1.00 | |
| **Average protein-coding gene length** | **base pairs** | | | | 3230 |
| | **amino acids** | | | | 365.8 |
| **BUSCO (viridiplantae db10)** | complete | singletons | duplicates | missing | fragmented |
| **transcripts** | 97.10% | 4.90% | 92.20% | 1.70% | 1.20% |
| **proteins** | 94.40% | 8.50% | 85.90% | 3.50% | 2.10% |

AED = Annotation Edit Distance (reference 49); BUSCO = Benchmarking Universal Single-Copy Orthologs (reference 43)

### Supplementary Table 4

| Gene | Species | Gene Number | Chr / Contig |
|---|---|---|---|
| DAM1 | *P. cerasus* | Pcer_004918-RA | chr1A |
| DAM2 | *P. cerasus* | Pcer_004919-RA | chr1A |
| DAM3 | *P. cerasus* | Pcer_004920-RA | chr1A |
| DAM4 | *P. cerasus* | Pcer_004921-RA | chr1A |
| DAM5 | *P. cerasus* | Pcer_004922-RA | chr1A |
| DAM6 | *P. cerasus* | Pcer_004923-RA | chr1A |
| DAM1 | *P. cerasus* | Pcer_010087-RA | chr1A' |
| DAM2 | *P. cerasus* | Pcer_010088-RA | chr1A' |
| DAM3 | *P. cerasus* | Pcer_010089-RA | chr1A' |
| DAM4 | *P. cerasus* | Pcer_010090-RA | chr1A' |
| DAM5 | *P. cerasus* | Pcer_010091-RA | chr1A' |
| DAM6 | *P. cerasus* | Pcer_010092-RA | chr1A' |
| DAM1 | *P. cerasus* | Pcer_015227-RA | chr1B |
| DAM2 | *P. cerasus* | Pcer_015228-RA | chr1B |
| DAM3 | *P. cerasus* | Pcer_015229-RA | chr1B |
| DAM4 | *P. cerasus* | Pcer_015230-RA | chr1B |
| DAM5 | *P. cerasus* | Pcer_015231-RA | chr1B |
| DAM6 | *P. cerasus* | Pcer_015232-RA | chr1B |
| DAM1 | *P. fruticosa* | Pfrut_002595-RA | frut_contig8 |
| DAM2 | *P. fruticosa* | Pfrut_002594-RA | frut_contig8 |
| DAM3 | *P. fruticosa* | Pfrut_002591-RA | frut_contig8 |
| DAM4 | *P. fruticosa* | Pfrut_002590-RA | frut_contig8 |
| DAM5 | *P. fruticosa* | Pfrut_002589-RA | frut_contig8 |
| DAM6 | *P. fruticosa* | Pfrut_002588-RA | frut_contig8 |
| DAM1 | *P. fruticosa* | Pfrut_003725-RA | frut_contig33 |
| DAM2 | *P. fruticosa* | Pfrut_003726-RA | frut_contig33 |
| DAM3 | *P. fruticosa* | Pfrut_003727-RA | frut_contig33 |
| DAM4 | *P. fruticosa* | Pfrut_003728-RA | frut_contig33 |
| DAM5 | *P. fruticosa* | Pfrut_003729-RA | frut_contig33 |
| DAM6 | *P. fruticosa* | Pfrut_003730-RA | frut_contig33 |
| **Gene** | **Species** | **NCBI identifier** | |
| DAM1 | *P. avium* | QYK20666.1 | |
| DAM2 | *P. avium* | QYK20667.1 | |
| DAM3 | *P. avium* | QYK20668.1 | |
| DAM4 | *P. avium* | QYK20669.1 | |
| DAM5 | *P. avium* | QYK20670.1 | |
| DAM6 | *P. avium* | QYK20671.1 | |
| SEP3 | *A. thaliana* | NP_001185081.1 | |

DAM (Dormancy Associated MADS-box) gene numbers in the ‘Montmorency’ and *Prunus fruticosa* assemblies, as well as the NCBI identifiers for the sequences from *P. avium* used in the phylogenetic analysis (Fig. 5b).

### Supplementary Table 5

| Gene | Species | Gene Number | Chr / Contig |
|---|---|---|---|
| *S*- RNase 36a | *P. cerasus* | Pcer_019010-RA | chr6A |
| SFB 36a | *P. cerasus* | Pcer_019009-RA | chr6A |
| *S*-RNase 35 | *P. cerasus* | Pcer_044499-RA | chr6A' |
| SFB 35 | *P. cerasus* | Pcer_044497-RA | chr6A' |
| *S*-RNase 13m | *P. cerasus* | Pcer_022874-RA | chr6B |
| SFB 13 | *P. cerasus* | Pcer_022871-RA | chr6B |
| *S*-RNase 6 | *P. cerasus* | Pcer_022887-RA | chr6B |
| SFB 6 | *P. cerasus* | Pcer_022886-RA | chr6B |
| *S*-RNase 36b | *P. fruticosa* | Pfrut_102374-RA | frut_contig53 |
| SFB 36b | *P. fruticosa* | Pfrut_102375-RA | frut_contig53 |
| **Gene** | **Species** | **NCBI identifier** | |
| *S*- RNase 36b | *P. cerasus* | EU042128.1 | |
| *S*- RNase 36b2 | *P. cerasus* | EU042129.1 | |
| *S*- RNase 36b3 | *P. cerasus* | EU042130.1 | |
| *S*- RNase 36a | *P. cerasus* | EU042127.1 | |
| *S*-RNase 35 | *P. cerasus* | EU054327.1 | |
| *S*-RNase 6 | *P. avium* | DQ385841.1 | |
| *S*-RNase 13m | *P. cerasus* | DQ385843.1 | |
| SFB 36b | *P. cerasus* | EU042132.1 | |
| SFB 36b2 | *P. cerasus* | EU042133.1 | |
| SFB 36b3 | *P. cerasus* | EU042134.1 | |
| SFB 35 | *P. cerasus* | EU054330.1 | |
| SFB 36a | *P. cerasus* | EU042131.1 | |
| SFB 13 | *P. avium* | DQ385844.1 | |

*S*-RNase and SFB genes, which together comprise the *S*-haplotype, identified in the 'Montmorency' and *P. fruticosa* assemblies; also, the NCBI identifiers for the sequences used as queries in the BLAST+ search.
